# Attachment Stimuli Trigger Widespread Synchrony across Multiple Brains

**DOI:** 10.1101/2023.02.10.527970

**Authors:** Ortal Shimon-Raz, Yaara Yeshurun, Adi Ulmer-Yaniv, Ayelet Levinkron, Roy Salomon, Ruth Feldman

## Abstract

Infant stimuli elicit widespread neural and behavioral response in human adults and such massive allocation of resources attests to the evolutionary significance of the primary attachment. Here, we examined whether attachment-related cues also trigger cross-brain concordance, generating greater neural uniformity among individuals. Post-partum mothers were imaged twice in oxytocin/placebo administration design and stimuli included four ecological videos; two of infant/mother alone (*Alone*) and two mother-infant dyadic contexts (*Social*). Theory-driven analysis measured cross-brain synchrony in preregistered nodes of the parental caregiving network (PCN), which integrates subcortical structures underpinning mammalian mothering with cortical areas implicated in simulation, mentalization, and emotion regulation, and data--driven analysis assessed brain-wide concordance using whole brain parcellation. Results demonstrated widespread cross-brain synchrony in both the PCN and across the neuroaxis, from primary sensory and somatosensory areas, through insular-cingulate regions, to temporal and prefrontal cortices. The *Social* context yielded significantly more cross-brain concordance, with PCN’s striatum, parahipporcampal gyrus, superior temporal sulcus, anterior cingulate cortex (ACC), and prefrontal cortex displaying cross-brain synchrony only to mother-infant social cues. Moment-by-moment fluctuations in mother-infant social synchrony, ranging from episodes of gaze aversion to rhythmically-coordinated positive bouts, were tracked online by cross-brain concordance in the pre-registered ACC. Findings indicate that social attachment stimuli, representing evolutionary-salient universal cues that require no verbal narrative for immediate comprehension, trigger substantial inter-brain concordance and suggest that the mother-infant bond, an icon standing at the heart of human civilization, may function to glue brains into a unified experience and bind humans into social groups.

**Significance Statement:** Infant stimuli elicit widespread neural response in human adults, attesting to their evolutionary significance, but do they also trigger cross-brain concordance and induce neural uniformity among perceivers? We measured cross-brain synchrony to ecological mother-infant videos. We employed theory-driven analysis, measuring cross-brain concordance in the parenting network, and data-driven analysis, assessing brain-wide concordance using whole-brain parcellation. Attachment cues triggered widespread cross-brain concordance in both the parenting network and across the neuroaxis. Moment-by-moment fluctuations in behavioral synchrony were tracked online by cross-brain variability in ACC. Attachment reminders bind humans’ brains into a unitary experience and stimuli characterized by social synchrony enhance neural similarity among participants, describing one mechanism by which attachment bonds provide the neural template for the consolidation of social groups.

## Introduction

Cross-brains synchrony research examines the correspondence in neural activations among individuals when exposed to the same dynamic stimulus, typically a movie or story (Hasson et al., 2004; Nastase et al., 2020; Visconti di Oleggio Castello et al., 2020). This neural synchronization taps the brain’s pre-wired response to the unfolding of events and pinpoints regions that curate such cross-brains resemblance. Studies on cross-brains mechanisms indicate that similarity in neural response increases when individuals share a narrative (Grall et al., 2021) or view emotionally-arousing movies (Nummenmaa et al., 2012); brains “tick together” when individuals understand a story in the same way (Yeshurun et al., 2017) or become emotionally engaged (Song et al., 2021). It has been theorized that processes of cross-brain synchrony played an important role in the evolution of human sociality, providing the foundation for human communication, collaborative abilities, and the capacity for empathy that sustain the formation of social groups and shared cultural meaning-systems (Hasson et al., 2012; Feldman, 2021a). As such, specifying the processes that underpin the convergent response of multiple brains to ongoing shared events may provide valuable insights into the co-evolution of the social brain and human cultural heritage.

Most cross-brain studies to date presented participants with some form of a narrative (Saalasti et al., 2019; Grall et al., 2021; Nastase et al., 2021) and, across studies, areas of the default mode network (DMN) were found to exhibit high cross-brain concordance when individuals are exposed to the unfolding of narrative and integrate it with internal memories or prior knowledge (Yeshurun et al., 2021). Other areas displaying high cross-brain synchrony are primary sensory regions, higher-order associative areas, such as the STS (Hasson et al., 2004), and prefrontal regions (Nguyen et al., 2019), and synchrony in these areas is thought to index the emergence of shared experiences that rely on the similarity of neural response. Cross-brain synchrony in subcortical regions, such as amygdala or basal ganglia, is less common and was found in response to emotionally-arousing musical stimuli with levels of inter-subject correlations (ISC), the main metric for cross-brain synchrony, linked with moment-by-moment rating of negative valence (Trost et al., 2015), suggesting that fluctuations in cross-brain correlations track ongoing variations in the emotional features of the stimulus. Multimodal stimuli of musical instrument learning were also found to elicit cross-brain synchrony in paralimbic regions, including the PHG and insula (Fasano et al., 2020), indicating that activity related to daily living experiences triggers cross-brain concordance in insular cortex.

Stimuli depicting the mother-infant attachment elicit a widespread response in the brains of mothers, fathers, and non-parents across the neuroaxis, including subcortical regions, such as the amygdala, VTA, nucleus accumbens, and PHG; paralimbic areas, including the insula and ACC; and cortical areas, particularly the mPFC (Swain, 2008; Parsons et al., 2013; Abraham et al., 2014; Shimon-Raz et al., 2021), and these areas cohere into the global “parental caregiving network” (PCN, see (Feldman, 2015, 2017). To trigger response in the PCN, studies have utilized stimuli of an infant alone (Ranote et al., 2004; Noriuchi et al., 2008; Strathearn et al., 2008; Parsons et al., 2017), a parent alone during infant-related daily activity (Abraham et al., 2017), or videos of the parent and infant together in various ecological contexts (Kuo et al., 2012; Musser et al., 2012). Yet, while studies have shown activations in nodes of the PCN to both infant/parent alone and parent-infant social stimuli, little empirical attention has been directed to cross-brain synchronization in response to attachment cues. As such, the main goal of the current study was to test whether stimuli representing the mother-infant attachment, a universal cue that requires no narrative for immediate comprehension, would elicit cross-brain synchronization in areas of the PCN and how elaborate is the neural concordance. Second, we wished to evaluate the effects of the social versus alone attachment context on cross-brain synchronization and test whether the degree of neural concordance in response to mother-infant dyadic stimuli (“social”) would be greater than to cues representing mother or child alone (“alone”). Finally, as oxytocin is a key modulator of the maternal brain in both humans (Numan, 2006; Norholt, 2020) and other mammals (Bosch and Neumann, 2012; Feldman and Bakermans-Kranenburg, 2017; Althammer et al., 2018), and OT administration impacts activity and connectivity in the PCN to attachment stimuli (Riem et al., 2011b; Bos et al., 2018; Shimon-Raz et al., 2021), we tested the effects of OT administration on cross-brain synchrony in response to four “social” and “alone” attachment stimuli. Two prior studies examined the effect of OT administration on cross-brain synchrony; an EEG study showed enhanced cross-brain synchrony under OT (Mu et al., 2016) and a functional magnetic resonance imaging (fMRI) study showed that OT administration modulated inter-brain correlations in the dorsal DMN and the precuneus network (Wu et al., 2022), and we thus examined the effects of OT on both the PCN and DMN.

Postpartum mothers, for whom daily mother-infant contexts are the most relevant, rewarding, and arousing (Kim et al., 2016; Parsons et al., 2017) were presented with four movies depicting daily mother-infant “social” and “alone” contexts. Utilizing a double-blind placebo-controlled oxytocin-administration crossover design, mothers’ brains were imaged twice a week apart. We used a dual analytic approach to examine cross-brain synchrony to attachment cues; “theory driven” and “data driven”. The “theory driven” approach was based on multiple imaging studies that pinpointed nodes of the PCN and these were pre-registered, along with the DMN, as our regions of interest (ROIs) and expected to show above-threshold cross-brain synchronization to attachment cues. The “data-driven” analysis used resting-state-based parcellation to examine inter-subject correlation across the entire brain (Shen et al., 2013). These methods combined a top-down with a bottom-up approach to define areas that exhibit cross-brain synchronization to attachment cues.

Three hypotheses were pre-registered. First, we hypothesized that a synchronized response, as measured by inter-subject correlation (ISC), in brain regions of the PCN and DMN would be found when mothers observe ecological videos of attachment contexts. As mother’s brain is selectively wired to social moments (Shimon-Raz et al., 2021), our second hypothesis contended that the social contexts would elicit greater synchronization in the PCN as compared to contexts of mother or infant alone. Finally, as OT is a key modulator of the maternal brain (Panksepp et al., 1997; Marlin and Froemke, 2017; Sanson and Bosch, 2022), we hypothesized that OT would modulate cross-brain synchronization levels in postpartum mothers. In addition to these pre-registered hypotheses, we also examined whether modulations in behavioral synchrony during the free play vignette will be tracked by modulations in mothers’ cross-brain synchronization. The ACC and insula play a key role in the neural representation of attachment (Ulmer-Yaniv et al., 2022), and we expected that synchronization fluctuations in the ACC and insula may track fluctuations in levels of mother-infant behavioral synchrony when mothers are exposed to attachment-related social cues.

## Materials and Methods

### Participants

The initial sample included thirty-five postpartum mothers who were recruited through advertisements in online parenting forums. Following recruitment, mothers underwent a brief phone screening for MRI scanning and postpartum depression using the Edinburgh Postnatal Depression Scale (EPDS) (Cox et al., 1987). Cutoff for joining the study was EPDS score of 8 and below (score above 9 indicates minor depression). Next, mothers were invited to a psychiatric clinic for a psychiatric evaluation prior to the scanning sessions. During this visit, mothers were interviewed using the Structured Clinical Interview for the DSM-IV (SCID) to assess current and past psychiatric disorders. None of the participants met criteria for a major or minor depressive episode during the perinatal period, 97% did not meet criteria for any diagnosable psychopathology during this time, and 86% did not meet criteria for any diagnosable psychopathology disorder during their lifetime.

All participants were married, cohabitated with the infant’s father, were of middle-or upper-class socioeconomic status, and completed at least some college.

Of the 35 participants, 3 did not complete a single scan (one due to medical problems and two due to claustrophobia). After examining the quality of the data, 4 mothers were excluded due to excessive head movement artifacts (movements ≥3 mm). In an additional participant we identified unexplained noise in the signal, detected by contrasting the visual conditions vs. rest. Three other mothers watched a different set of stimuli and were unable to enter the ISC analysis. These 11 subjects were removed before analysis of the experimental effects.

The final sample used for the analysis included 24 mothers (mean age = 29.62 years, SD = 4.9; EPDS mean score = 3.12, SD = 2.50) of 4-7-month-old infants (mean age = 5.58 months, SD = 1.37) and each mother underwent scanning twice (48 scans). A sample size of 24 participants has been shown in previous studies to be sufficient for assessing neural responses to naturalistic stimuli (Ames et al., 2015; Yeshurun et al., 2017) and for power analyses of inter-subject correlation, the analytic method in the current study (Pajula and Tohka, 2016).

The study was approved by the Helsinki committee of the Sourasky Medical Center, Tel Aviv (Ethical approval no. 0161-14-TLV). All participants signed an informed consent. Subjects received a gift certificate of 700 NIS (~200 USD) for their participation in all 4 phases of the study (diagnosis, home visit, and two imaging sessions).

### Stimuli and experimental design

Following psychiatric evaluation, the study included three sessions for each mother. In the first, families were visited at home and episodes of mother-infant free-play interaction, infant alone, mother alone and breastfeeding were videotaped. In addition, mothers completed self-report measures.

In the second and the third sessions, mothers participated in fMRI brain scanning at the Tel-Aviv Sourasky Medical Center. Before each scan mothers received 24 IU of placebo or oxytocin intranasally in a randomized, placebo-controlled, double-blind, two-period crossover design. On average, 14 days elapsed between the two scans (SD = 9.30, mode = 7, median = 7), that were both scheduled for the morning hours (07:30–12:00).

The mother-infant context paradigm and fMRI sequence began about 30 minutes after intranasal placebo (PBO)/ Oxytocin (OT) administration. During the scans, each mother was presented with 8 naturalistic films of 120 seconds each: four videos of the same unfamiliar mother and her infant, and four individually tailored matched videos of the mother herself and her own infant. In this paper, only the neural response to the standard-unfamiliar mother and infant videos was analyzed and is therefore referred as our two stimuli. The *Alone* stimulus included 2 videos, one of the infant alone (in the bouncer, playing with a toy) and one of the mother alone (sitting, folding laundry), while the *Social* stimulus included 2 videos of the mother and the infant together, one during a free-play (playing together while the baby is in the bouncer) and one during breastfeeding. Between videos, a fixation of a black cross over a grey background was presented. Fixation duration alternated between 15 and 18 seconds. The order of the videos was counterbalanced across participants and scans. While in the scanner, mothers were asked to watch the movies attentively. Video clips were played using VLC media-player (version 2.2 for Windows, VideoLAN, France). Study procedure and fMRI paradigm are presented in Fig. 1A.

**Fig. 1.**
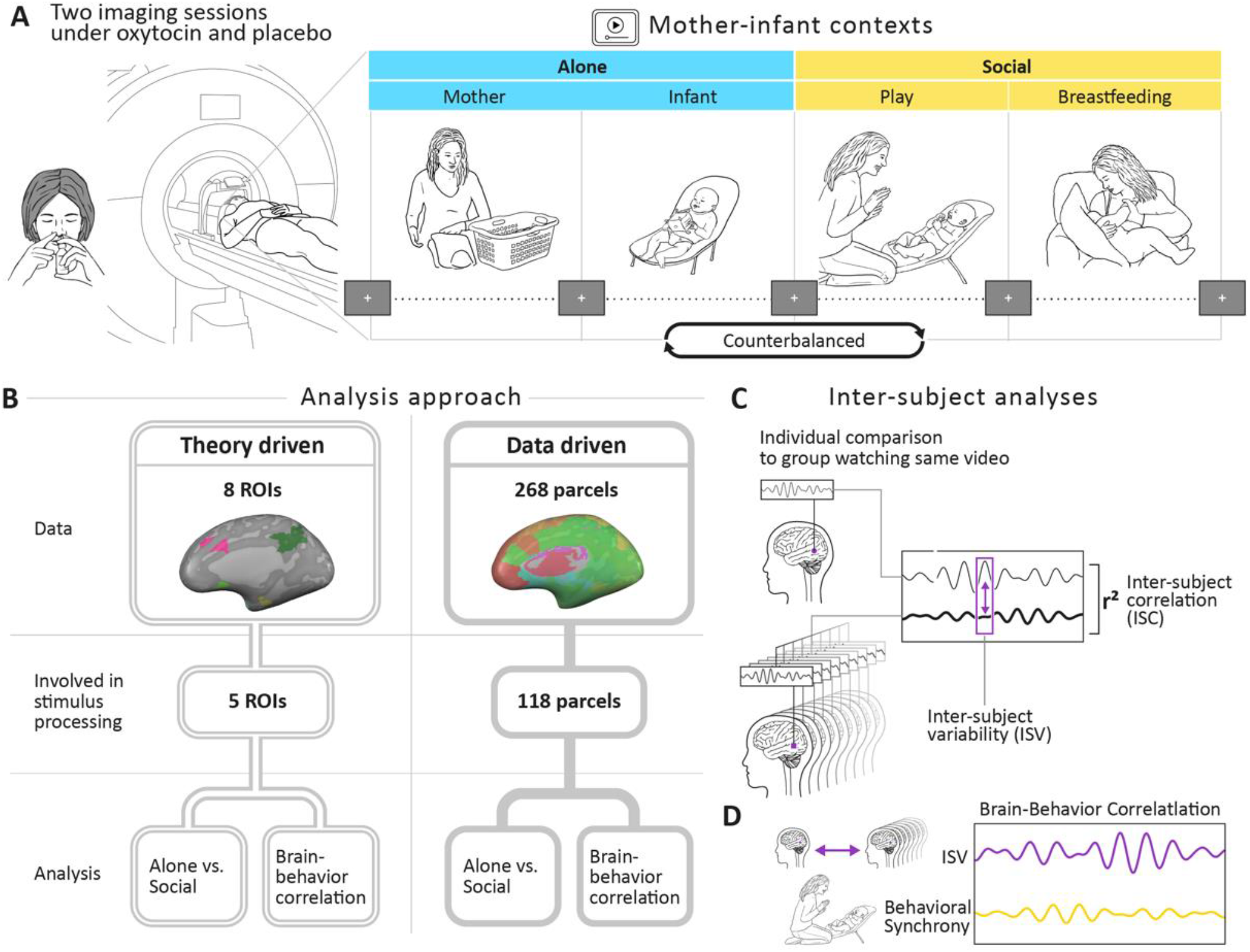
Research plan. **A**. Experimental procedure and paradigm. Post-partum mothers were imaged twice after oxytocin/placebo administration in a randomized, placebo-controlled, double-blind, crossover design. On average two weeks elapsed between scans. While in the scanner participants observed 4 daily ecological video vignettes showing a standard mother and an infant apart and together. The *Alone* context condition (in blue) included two videos of the mother alone while folding infant laundry and of the baby alone sitting in the bassinet while playing with a toy. In the *Social* context condition (in yellow), two videos showed the mother and the infant together while engaged in free-play interaction and during a breastfeeding episode. Videos lasted 2 minutes each and were previewed by rest with fixation period of 1 min. A rest with fixation periods of alternately 15-18 seconds was presented between clips. Order of videos was counterbalanced between the two scans. Bayesian ANOVA results of two conditions are presented in Fig. 1-1 & Fig. 1-2. **B**. Two approaches were used to examine differences in inter-subject correlation (ISC): a theory-driven approach focused on 8 preregistered ROIs: 7 within the PCN and the DMN (Fig. 1-3.), and a data-driven approach that tested the entire brain using Shen’s (2013) atlas that consist of 268 parcels. In the two approaches we first calculated ISC value for each brain area and compared it to ISC threshold. Regions showing ISC above threshold were defined as involved in stimulus processing. In these areas we explored differences between the ISC in the Alone and the Social conditions, and the correlation between the participants’ moment-by-moment neural similarity and the continuous level of mother-infant behavioral synchrony in the free-play interaction. **C**. ISC measures the neural similarity across participants by correlating the time course of one participant with the averaged time course of all other participants in the same brain region. Inter Subject Variability (ISV) is a complimentary measure that reflects the neural variability of the group in a certain timepoint. It was calculated using Euclidean Distance (ED), which is the difference in the BOLD signal at a certain time between a subject and the rest of the group’s mean. **D**. Brain-behavior correlation was examined in response to the free-play interaction video. In each TR mother-infant behavioral synchrony was evaluated using a micro-coding system and correlated with the continuous time course of the ISV. ISV score was transformed to z score and multiplied by −1 in order that higher values will represent greater neural/behavioral synchrony. For detailed description of the behavioral coding see Fig.1-4. Abbreviations: PCN, parental caregiving network; DMN, default mode network.

### MRI acquisition

Magnetic Resonance Imaging (MRI) data was collected using a 3T scanner (SIEMENS MAGNETOM Prisma syngo MR D13D, Erlangen, Germany) located at the Tel Aviv Sourasky Medical Center. Scanning was conducted with a 20-channel head coil for parallel imaging. Head motion was minimized by padding the head with cushions, and participants were asked to lie still during the scan. High resolution anatomical T1 images were acquired using magnetization prepared rapid gradient echo (MPRAGE) sequence: TR = 1860 ms, TE = 2.74 ms, FoV = 256 mm, Voxel size = 1x1x1 mm, flip angle = 8°. Following, functional images were acquired using EPI gradient echo sequence. TR = 3000 ms, TE = 35 ms, 44 slices, slice thickness = 3 mm, FOV = 220 mm, Voxel size = 2.3x2.3x3 mm^3^, flip angle = 90°. In total 381 volumes were acquired over the course of the “Context” paradigm. Visual stimuli were displayed to subjects inside the scanner, using a projector (Epson PowerLite 74C, resolution = 1024 × 768), and were back-projected onto a screen mounted above subjects’ heads, and seen by the subjects via an angled mirror. The stimuli were delivered using “Presentation” software (www.neurobs.com).

### Oxytocin administration

Mothers were asked to self-administer 24 IU of either oxytocin (Syntocinon Nasal spray, Novartis, Basel, Switzerland; three puffs per nostril, each containing 4 IU) or placebo prior to scanning. The placebo was custom designed by a commercial compounding pharmacy to match drug solution without the active ingredient. The same type of standard pump-actuated nasal spray was used for both treatments.

### Behavioral data analysis

#### Micro-coding of social synchrony during mother-infant free-play condition

To track the variability in the degree of behavioral synchrony during the free-play film, we applied the *Parent-Infant Synchrony* (Feldman and Eidelman, 2007) coding scheme to the free play session, consistent with our previous research (Feldman and Eidelman, 2003, 2007; Feldman et al., 2004) including fMRI studies (Atzil et al., 2011; Shimon-Raz et al., 2021). Micro-coding was performed by a trained coder on a computerized system (Mangold- Interact, Arnstorf, Germany, RRID: SCR_019254) in 3 seconds frames. Three non-verbal aspects of social behavior was coded for mother and infant separately; *Affect* (very positive, positive, neutral, negative withdrawn, negative/upset, uncodable), *Gaze* (infant: to mother, to object, joint gaze to object, to environment, sleepy/drowsy, aversion, uncodable; mother: to infant’s face, to infant’s body, to object, joint gaze to object, to environment, aversion, uncodable), and *Vocalization* (infant: cooing, fussing, laughing, crying, no vocalization, uncodable; mother: motherese, adult directed speech to infant, adult speech to another adult, laugh, no vocalization, uncodable). Synchrony was defined, consistent with our prior research (Feldman and Eidelman, 2007), by conditional probabilities (infant in state A given mother in state A), indicating episodes when the mother and the infant were both in social gaze, positive vocalizations, and shared positive affect (Feldman and Eidelman, 2007; Granat et al., 2017). The degree of synchrony was graded on a scale from 1 to 15. The lowest level of synchrony (coded as level 1) involved mother looking at infant with neutral affect and no vocalization and infant gaze-averting and expressing no positive affect or vocalization. The highest level of synchrony (coded as level 15) involved mother looking at infant, expressing positive affect, and vocalizing while infant looked at mother’s face, was positive, and laughed. A detailed description of the 15 levels of synchrony is provided in Supplementary Table 1.

### MRI data Analysis

#### Data preprocessing

Data preprocessing and data analysis were conducted using BrainVoyager QX software package 20.6 (Brain Innovation, Maastricht, The Netherlands, RRID: SCR_013057) (Goebel et al., 2006). The first 3 functional volumes, before signal stabilization, were automatically discarded by the scanner to allow for T1 equilibrium. Preprocessing of functional scans included 3D motion correction, slice scan time correction, spatial smoothing by a full width at half maximum (FWHM) 6-mm Gaussian kernel, and temporal high-pass filtering. The functional images were then manually aligned and co-registered with 2D anatomical images and incorporated into the 3D datasets through trilinear interpolation. The complete dataset was normalized into Montreal Neurological Institute (MNI) space (Evans et al., 1994).

The scanning sessions resulted in 40-TR long recordings of blood oxygenation level dependent (BOLD) signal intensity per film, from which the first two TRs (equivalent to 6 seconds) were excluded, to consider hemodynamic response. ROI and parcel-specific BOLD time-course were produced for each subject and video, by averaging the time-series of all voxels in the same area. Z scores were calculated for each time series separately, giving each TR a normalized value representing signal intensity

#### Regions-of-Interest Preregistration and Analysis

ROI analysis was conducted on eight preregistered bilaterally defined ROIs (https://osf.io/bmp43?view_only=0ca4cb28ef2c4b4092a44a1c047c9242), including the amygdala, anterior cingulate cortex (ACC), anterior insula, hippocampus/ parahippocampal gyrus, temporal pole, ventral tegmental area (VTA), nucleus accumbens (NAcc), (all together defined as the “maternal caregiving network”), and the default mode network (DMN). ROIs were selected a-priori, based on theory and literary meta-reviews (Abraham et al., 2016; Lindquist et al., 2016), and on a pilot study of 4 subjects that completed similar paradigm and were not included in the current study. ROIs were defined functionally and anatomically, verified and validated by human brain database platforms: Talairach Daemon (Lancaster et al., 2000) and Neurosynth (Yarkoni et al., 2011), registered at the Open Science Framework prior to data analysis (OSF, n.d.) and transformed into MNI space.

Time-courses were extracted from ROIs, and ISC for each ROI was calculated. After thresholding, 5 ROIs, 4 of the PCN and the DMN were found to be involved in stimulus processing and analyzed with two separate repeated measures ANOVA. In the PCN a 4×2×2 (*ROI_PCN_* × *Context* × *PBO-OT*) was computed and in the DMN a 2×2 (*Context* × *PBO-OT)* was calculated. Thus, allowing to investigate main effects of stimulus type, oxytocin administration, and their interactions. In order to further examine the origin of main effects and interactions, simple effect analyses, and FDR corrected post hoc tests were conducted.

#### Data Driven Whole-Brain Analysis

In the current study, we used ISC to identify regions that were involved in processing the video-clips. We conducted both whole-brain analysis using Shen’s parcellation (Shen et al., 2013) and ROI analysis on predefined areas. Shen’s whole-brain parcellation defines 268 parcels based on resting-state fMRI data that yielded nodes with coherent internal time-courses (Shen et al., 2013). In the current study parcels were labeled with their serial number, which ranged from 1 to 268 in the atlas, as well as their location association in meta-analysis maps in the Neurosynth human brain database platform.

ISC and *p* values were calculated for the 4 conditions (*Alone/Social* × *PBO/OT*) in each parcel. Next, ISC threshold and FDR corrected *p* values were calculated for each condition. Parcels which passed this thresholding in at least one condition were defined as involved in stimulus processing. Following, a two factor (*Context* × *PBO-OT*) repeated measures ANOVA was performed and yielded 2 main effects and an interaction effect for each parcel.

#### Inter subject correlation

To test our preregistered hypothesis that *Social* videos would yield greater neural similarity among subjects than *Alone* videos, we compared both condition’s inter-subject correlation (ISC), a measure of neural response coherence across individuals over the time-course of a stimulus.

We calculated an ISC score for a given brain region (ROI/parcel) by correlating each participant’s time course with the averaged time course of the other participants using Pearson’s correlation. This procedure resulted in 24 ISC values (one per participant) which were averaged to obtain one ISC value per region. Higher ISC imply more synchronized brain responses to the stimuli.

Analysis of effects within a 2 factors Bayesian repeated measures ANOVA (*Infant/Mother* alone movie × *PBO-OT*) showed moderate evidence for the absence of difference between the mother alone and the infant alone movies (*BF*_10_ = 0.25, BF_incl_ = 0.18), and similar analysis that was computed and revealed moderate evidence against the difference between the play and the feeding movies (BF_10_ = 0.341, BF_incl_ = 0.246) (Supplementary Tables 2,3). For testing the response to *Alone* versus *Social* stimuli, time-courses sampled during the presentation of the infant alone video and the mother alone video, 38-TRs each, were concatenated into a 76-TRs long sequence of an *Alone* condition, while time-courses of the *breastfeeding* and the mother-infant *free-play* videos were concatenated into a *Social* condition sequence. Next, the two 76-TR sequences had their ISC coefficient calculated separately.

The structure of a subject *w*’s joined time-series at a certain ROI or parcel

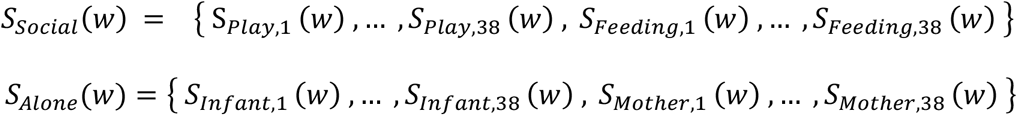

ISC scores significance was tested using bootstrapping. A resampling procedure was applied to each of the *Alone* and *Social* sequences for PBO/OT separately. Fast Fourier Transform (FFT) was used to separate each sequence into independent components while preserving the power spectrum of the signal. Then, 10,000 randomized phase permutations of the sequences were assembled using inverse-FFT. By calculating each of the new time-courses’ ISC, we achieved a null distribution of 10,000 ISC scores.

In each condition, given an area (either ROI or parcel) with an average ISC score of 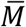, the *p* value for that area was determined by the proportion of ISC scores greater than 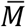 in the null distribution. Adding one to the count of said values ensures *p* is a non-zero probability.

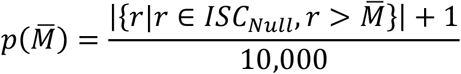

Critical *p*-values were calculated using Benjamini-Hochberg’s correction for controlling the false-discovery rate (Benjamini & Hochberg, 1995):

– Sort the obtained *p*-values: {*p*_1_, …, *p*_*m*_} (*p*_*i*_ denotes the *i*-smallest *p*-value).
– For *i* = {1, …, *m*}, calculate 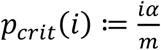 (where *m* is the number of comparisons and *α* is the probability threshold; in our case, *m* = 2 × 8 = 16, *α* = 0.05).
– Let *k* be the largest *i* ∈ [1, *m*] such that *p*_*i*_ ≤ *p*_*crit*_(*i*). Then the critical *p*-value is *p*_*crit*_(*k*).

To narrow down all tested regions to those which yielded the strongest coherence, an additional (more selective) threshold was applied. This threshold was the ISC score of the upper 5th percentile of the null distribution of each condition. Brain areas which passed that threshold, were defined as involved in stimulus processing and continued to ANOVA, as described later in the Results section.

#### Inter Subject Variability

Inter Subject Variability (ISV) is a measure of the magnitude of dispersion in ISC scores within the group at a certain time. The dispersion was determined using Euclidean Distance (ED), which is the difference in the BOLD signal magnitude at a certain time between a subject and the rest of the group’s mean. The ED of subject *s* at TR *t* can be calculated by the expression:

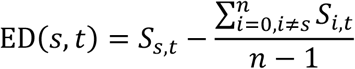

Where *n* is the number of subjects (24) and *S*_*i*,*t*_ is the value of the *t*^th^ sample in the normalized time-course of subject *i*.

ISV was calculated for the *Social* free-play condition in each ROI and parcel that were involved in stimulus processing (pass the ISC threshold) by averaging the squared ED values of each TR (between 24 subjects), then dividing by the greatest ED among all TRs in the time series corresponding to the *free-play* film.

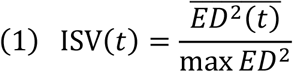

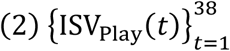

In order to quantify the similarity in responses instead of dispersion, we applied the following transformation for the resulting ISV time-series (Equation 2):

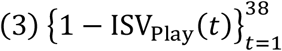

Next, Pearson’s *r* was calculated between the series of 38 average Z-scores (one for each TR) in the ISV complement time series (described in Equation 3), and the series of behavioral synchrony Z scores at each TR in the *free-play* video.

### Statistical analysis

For statistical analysis we used JASP (Version 0.9.2.0 for Windows, JASP Team, 2018, RRID: SCR_015823), SPSS (SPSS statistics Version 25.0, IBM Corp. Armonk, NY) and R software (Version 3.5.3, R Core Team, 2017, Vienna, Austria, RRID: SCR_019096). For all ISC calculations we used Matlab (Version 2021b, MathWorks Inc, Natick, MA).

## Results

### Brain regions showing cross-brain synchrony to attachment reminders

Consistent with our first preregistered hypothesis, daily ecological mother-infant contexts induced above threshold inter-subject brain synchronization in the parental caregiving network (PCN) and in the DMN. Four out of the 7 preregistered ROIs of the parental caregiving network as well as the DMN showed high ISC for at least one context under PBO or OT (all *p* values<.001). The ACC and the insula responded exclusively to the *Social* context videos (and not to the *Alone* videos), the PHG responded to the *Alone* context, and the NAcc and the DMN were involved in both contexts processing (Fig. 2A).

**Fig. 2.**
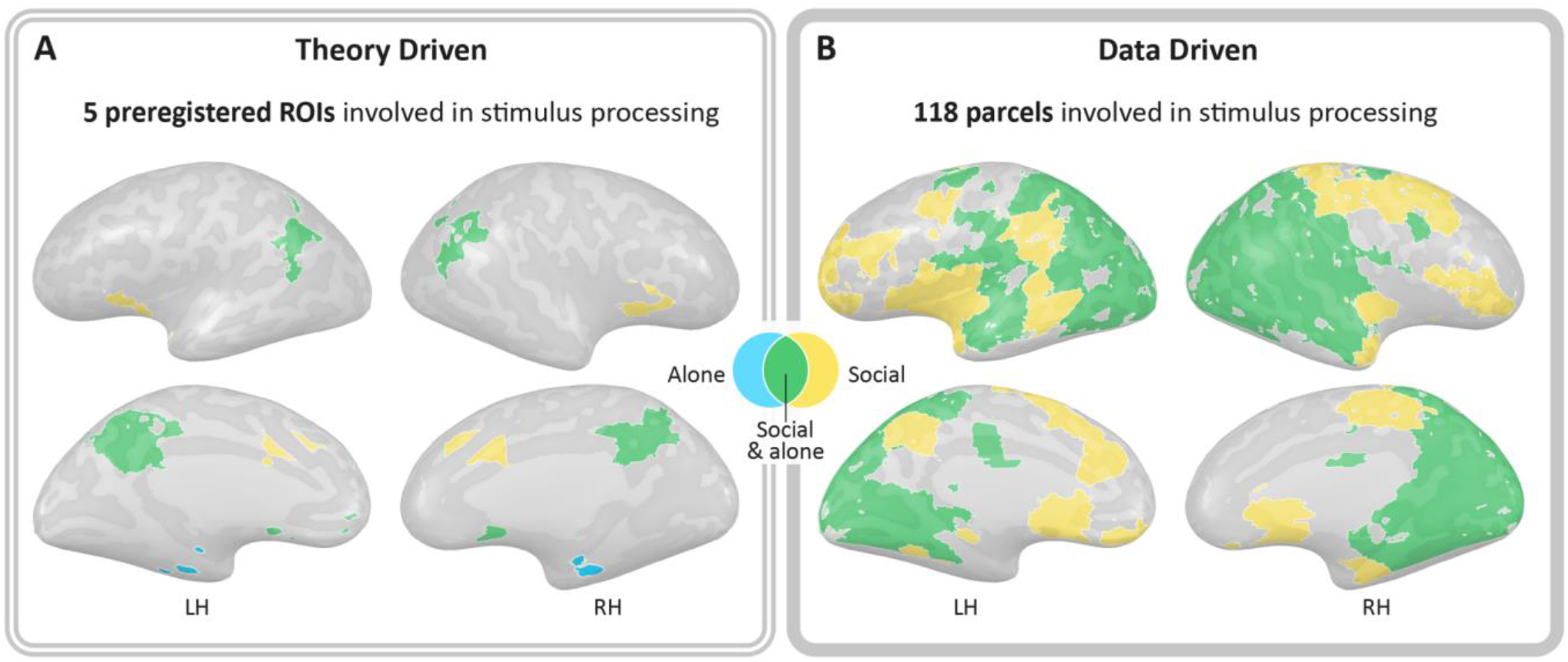
Figures represent brain regions involved in stimuli processing (significant ISC>ISC threshold, FDR corrected). In yellow-areas responded to the *Social* context, in blue-aeras responded to the *Alone* context, in green-areas responded to both, *Social* and *Alone* stimuli. **A**. Examination of the 8 preregistered ROIs revealed 5 areas that yielded high ISC including 4 areas of the PCN and the DMN. The preregistered insula and ACC responded to the *Social* context with high ISC, the PHG responded to the *Alone* context videos, the NAcc and the DMN responded to both conditions. **B**. 118 of 268 parcels showed high ISC in response to the stimuli (Fig.2-1). Along these, areas ranging from the subcortical striatum, insula, ACC, NAcc, PHG, to the STS, TPJ, premotor, visual, auditory, and somatosensory cortices, the OFC and prefrontal areas. 75 parcels responded to both stimuli. 42 responded exclusively to the *Social* condition and 1 parcel, in the cerebellum, responded only to the *Alone* condition. This highlights extensive similarity in neural response to attachment related stimuli particularly to the *Social* context. Note that similar areas were elicited in both analysis approaches. ISC thresholds are presented in Fig2-2, 2-3. Abbreviations: ISC, inter subject correlation; PCN, parental caregiving network; DMN, default mode network; ACC, Anterior cingulate; PHG, parahippocampal gyrus; NAcc, nucleus accumbens; STS, superior temporal sulcus; TPJ, temporoparietal junction; OFC, orbitofrontal cortex.

Furthermore, whole brain ISC revealed 118 parcels that were involved in processing the *Alone* and/or *Social* contexts videos, including areas in the PCN and the DMN. Out of these parcels, 75 parcels were involved in processing of both *Social* and *Alone* conditions. These regions included areas in the PFC, posterior insula, hippocampus, parahippocampal gyrus, cuneus and precuneus, fusiform gyrus, temporoparietal junction (TPJ), inferior parietal, premotor, visual, auditory and somatosensory cortices, and areas in the cerebellum (Fig. 2B). Forty-two parcels were only involved in processing the *Social* context, including areas in the PFC, orbitofrontal, superior temporal and premotor cortices, as well as subcortical striatum and the NACC, insula, ACC and hippocampus; and one parcel in the cerebellum was only involved in processing the *Alone* condition. For the full list of brain areas with significant ISC and ISC thresholds see Supplementary Table 5 and Table 6.

### *Social vs. Alone* contexts

The second preregistered hypothesis suggested increased ISC in the PCN in response to videos depicting mother-infant contexts compared to mother alone and infant alone videos.

For the 4 ROIs involved in stimulus processing within the PCN (ACC, insula, PHG& NAcc), a repeated measures ANOVA (*ROI* × *Context* × *PBO-OT*) revealed significant main effect of *Context*. The *Social* context elicited higher ISC compared to the *Alone* context [F(1,23) =10.96, p<0.01, Eta^2^=0.32], (Mean_*Social*_ = 0.14, SE_*Social*_=0.07 ; Mean_*Alone*_ = 0.08, SE_*Alone*_=0.06). Additionally, a significant main effect for *ROI* was found with the NAcc yielding higher ISC compared to the PHG [t(23)=3.28, p_bonf_= 0.02; Mean_NAcc_ = 0.15, SD_NAcc=_0.09 ; Mean_PHG_ = 0.07, SD_PHG_=0.06). Interaction effects were not significant (Supplementary Table 7). Post hoc repeated measures ANOVA (*Context* × *PBO-OT*) conducted in each of the ROIs revealed significant *Context* main effect in the insula and in the PHG, both driven by greater ISC in response to the *Social* context compared to the *Alone* context (Mean_*Social-Insula*_= 0.15, SD _*Social-Insula*_=0.08, Mean_*Alone-Insula*_= 0.07, SD_*Alone-Insula*_=0.09; Mean_*Social-PHG*_= 0.12, SD_*Social-PHG*_=0.11, Mean_*Alone-PHG*_=0.02, SD_*Alone-PHG*_=0.08) (Fig. 3A). No such effect was found in the ACC and the NAcc. In the DMN, which was also involved in stimulus processing, 2 factorial repeated measures ANOVA (*Context* × *PBO-OT*) didn’t reveal any significant effects (Supplementary Table 8).

**Fig. 3.**
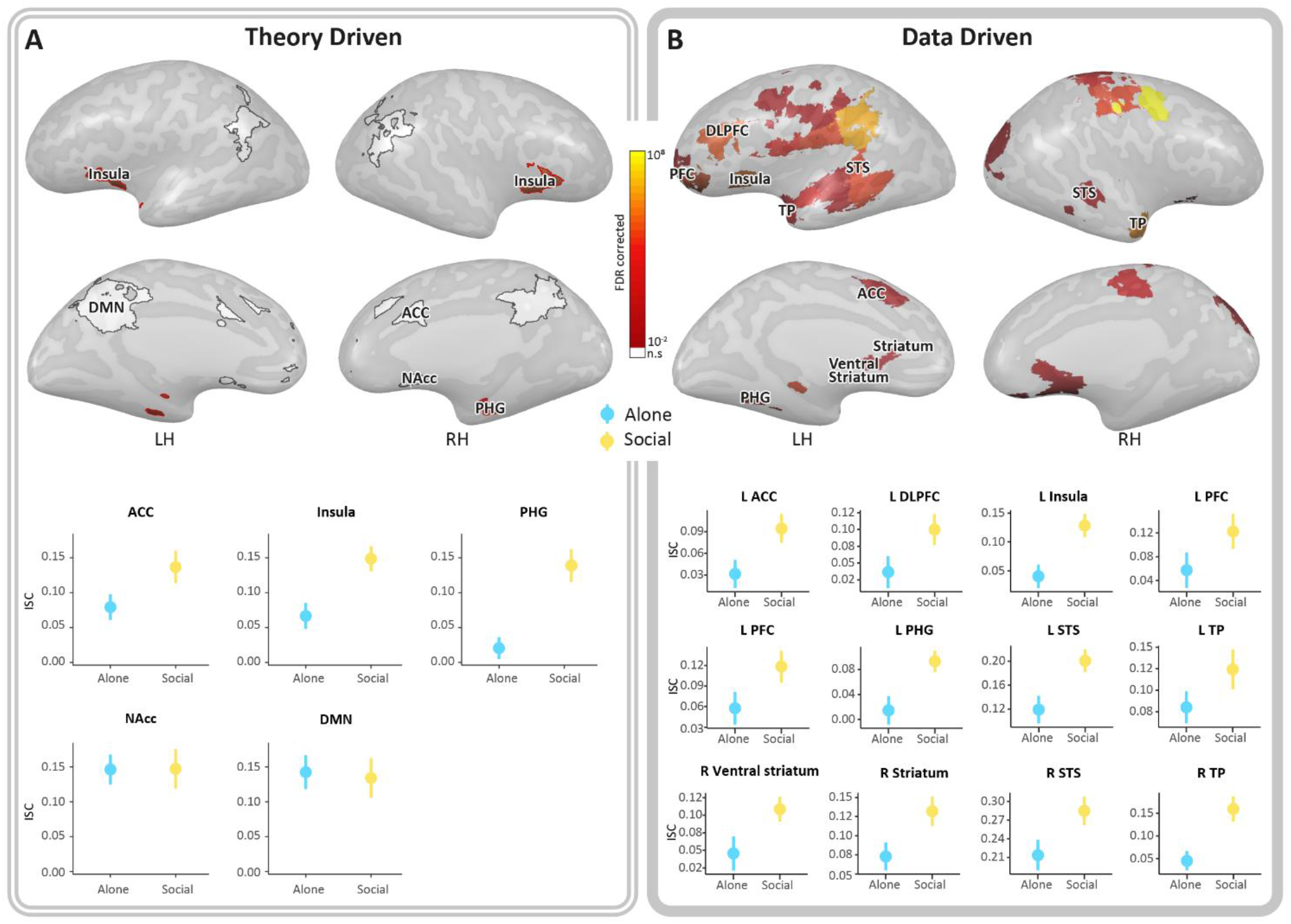
Significant main effect of *Context* was driven by greater ISC under the *Social* condition compared to the *Alone* condition. All results are FDR corrected. **A**. A repeated measures ANOVA (*ROI* × *Context* × *PBO-OT*) revealed significant main effect of *Context* that was driven by higher ISC under the *Social* context compared to the *Solitary* context in the preregistered insula and PHG (in red) (Fig. 3-1 & 3-2). ROIs in white color represent areas that responded to the stimuli but did not show such main effect. All graphs show ISC means in the *Social* (yellow) and in the *Alone* (blue) conditions. Bars depict standard error of the mean. **B**. Whole brain results of 2×2 repeated measures ANOVA (*Context* × *PBO-OT*) conducted for each of the 118 parcels involved in stimulus processing, revealed 27 parcels in which neural similarity was higher under the *social* condition (Fig.3-3). Within them 12 areas of the PCN that overlap with our ROIs and with Neurosynth map of the term “social” (Fig. 3-4). In the left hemisphere: ACC, Insula, DLPFC, PFC, PHG, STS, TP, and in the Right hemisphere: Striatum and VS, STS, TP. The level of significance of each region is indicated by a color scale ranging from red to yellow. Additional areas found include the OFC, motor, auditory and somatosensory cortices whose graphs are shown in Fig. 3-5. Abbreviations: PBO, placebo; OT, Oxytocin; PHG, parahippocampal gyrus; ACC, Anterior cingulate; TP, Temporal pole; VTA, ventral tegmental area; NAcc, Nucleus accumbens; DMN, default mode network; DLPFC, dorsolateral prefrontal cortex; PFC, prefrontal cortex; STS, superior temporal sulcus; TP, temporal pole; VS, Ventral striatum; OFC, orbitofrontal cortex. RH, right hemisphere; LH, left hemisphere.

The second hypothesis was also supported by data-driven analyses, which identified PCN regions and “social brain” areas with higher ISC in the social context versus the Alone context. A 2×2 repeated measures ANOVA (*Context* × *PBO-OT*) was performed for each of the 118 parcels involved in stimulus processing. In 27 parcels, a significant *Context* FDR-corrected effect for was found, with the *Social* context producing higher ISC compared to the *Alone* context (Fig. 3B). The ACC, insula, Superior temporal and parahippocampal cortex, orbitofrontal cortex, PFC, areas in the striatum, motor, and premotor cortices were among those parcels (Supplementary Table 9). Note that Results in the insula and PHG are consistent in both research approaches.

Regarding the third preregistered hypothesis which proposed enhanced ISC under OT compared to PBO, we found no significant differences between PBO and OT, neither in data, nor in theory driven analyses.

### Cross-brain synchrony tracks moment-by-moment variations in behavioral synchrony

Finally, we examined whether moments of behavioral synchrony induced greater neural synchrony between subjects, compared to non-synchronous moments. This was tested in brain areas that were involved only in the *Social* context processing under PBO or OT: 2 ROIs (ACC and Insula) and 42 parcels.

In the preregistered ACC, moments of greater mother-infant behavioral synchrony were associated with greater cross-brains synchrony between participants (*r_p_* = 0.445, *p* = 0.005) (Fig. 4A).

**Fig. 4.**
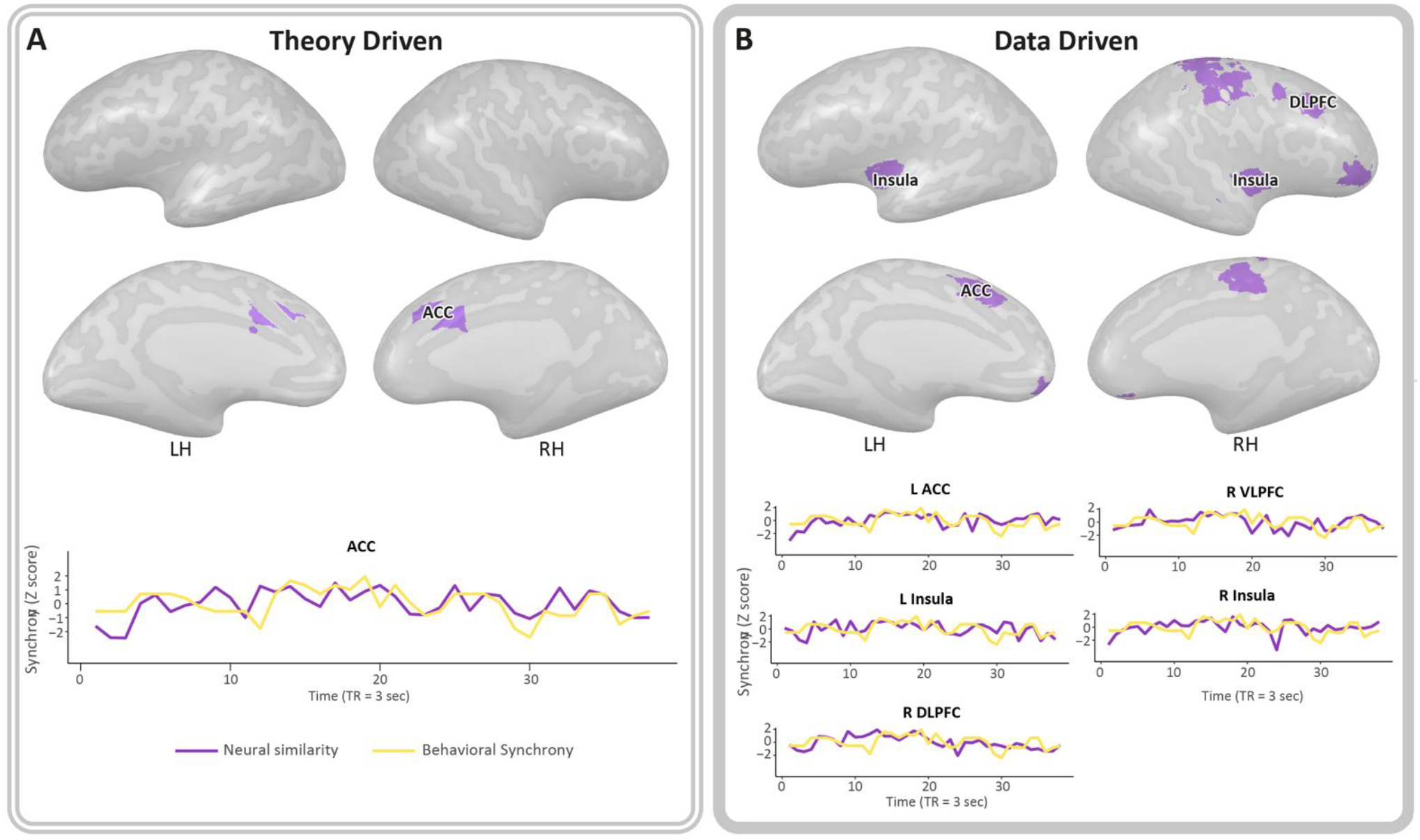
Brain-behavior correlations. The figure depicts brain areas where the level of cross-brain synchrony was correlated with mother-infant behavioral coordination in the free-play video that was viewed. **A**. Significant Pearson’s correlation showed that in the preregistered ACC moments of greater behavioral synchrony were related to greater neural similarity between subjects (non-significant results are in Fig.4-1). This result remains significant after multiple comparisons correction. The purple line in the graph represents neural similarity calculated by z transformation of the 1-ISV score; The yellow line represents the level of mother-infant behavioral synchronization based on micro-coding of the mother’s and baby’s gaze, affect and touch in each TR. **B.** Data driven analyses of Pearson’s correlations identified 11 parcels who showed significant positive brain-behavior correlation that did not survive correction for 42 comparisons. Among these, areas of the PCN: the left ACC, which overlap with the preregistered ACC, right and left insula, right VLPFC and right DLPFC. Their graphs are shown in the figure. Other areas that were found include the motor, primary motor, and somatosensory cortices, left and right OFC and the cerebellum (Fig. 4-2). Abbreviations: ISV, inter subject variability; ACC, Anterior cingulate; DLPFC, dorsolateral prefrontal cortex; VLPFC, ventrolateral prefrontal cortex; OFC, orbitofrontal cortex. RH, right hemisphere; LH, left hemisphere.

Similar findings were revealed in 11 parcels, in which more synchronous behavior yielded more synchronous brain activity among participants (Fig. 4B). This was found in areas of the “social brain” (i.e. OFC) and the PCN: right ACC (rp=0.42, p=0.009) left insula (rp=0.41, p=0.01), right VLPFC (rp=0.38, p=0.019) under PBO and OFC (right: rp=0.41, p=0.01; left: rp=0.35, p=0.03), right DLPFC (rp=0.36, p=0.02) and the right insula (rp=0.35, p=0.03), as well as in regions within the motor cortex (rp=0.43,p=0.007; rp=0.35,p=0.03), somatosensory cortex (rp=0.35,p=0.03) and in the cerebellum (rp=0.38,p=0.02). However, data driven results did not remain significant after FDR correction due to multiple comparisons.

## Discussion

The mother-infant bond is a defining feature of mammals, sustaining safety, growth, and adaptation of the young, and stimuli representing the primary attachment trigger a powerful response in the brain of human adults (Swain, 2011), as well as in the brain of other mammals (Vom Saal, 1985; Lonstein and De Vries, 2000; Novakov and Fleming, 2005; Olazábal and Young, 2006). While multiple studies described the brain areas activated in response to attachment reminders (Swain, 2011; Swain et al., 2014; Rigo et al., 2019), our results are the first to show substantial cross-brain synchronization to dynamic mother-infant stimuli. We found that large portions of the brain, in fact 44% of the parcels measured, activate in tandem to the presentation of attachment stimuli, indicating that bonding-related cues reduce variability among individual brains and enhance their synchronous, uniform response. Areas of cross-brain concordance were widespread, from lower-level visual, auditory, and somatosensory cortices, to subcortical limbic and paralimbic regions, to cortical prefrontal and orbitofrontal areas. Representation of the primary attachment, therefore, not only elicits a wide-spread neural response but also generates significant uniformity among multiple brains

Our findings indicate that while cross-brain similarity in largely-distributed networks ensue any reminder of the primary attachment, the *social* context representing the mother-infant bond triggers significantly greater neural concordance. In fact, 27 parcels (23% of the “synchronized parcels”), including regions of the PCN as well as sensorimotor, visual, auditory, and higher-level regions, activated synchronously to the presentation of mother-infant *social* stimuli, not to the *alone* contexts. Our results demonstrate not only cross-brain resemblance in most areas of the PCN, but also cross-brain synchronization in regions implicated in sensory processing and integration, mentalization, association, and higher-order valuation. Furthermore, our findings show for the first time that moment-by-moment variability in the magnitude of cross-brain synchronization tracks online fluctuations in levels of mother-infant behavioral synchrony and pinpoints the ACC in this cross-brains- behavior linkage. This suggests that mother-infant synchrony, a core human-specific behavioral mechanism that sustains online physiological (Feldman et al., 2011) and neural (Endevelt-Shapira et al., 2021) synchrony between mother and child during social moments and shapes children’s social-emotional and neural outcomes up till adulthood (Feldman, 2016, 2017, 2021b; Ulmer Yaniv et al., 2021), also functions to build synchrony across multiple brains. The behavioral building-blocks of social synchrony – gaze, affect, and vocalizations – are expressed between all mothers and infants in our species and, across cultures, assume a repetitive-rhythmic, highly coordinated expression. It has been suggested that through mechanisms of *biobehavioral synchrony*, which integrate the online coordination of physiological processes and social signals, mothers usher infants into the social world (Feldman, 2021a). The current findings raise the possibility that such moments not only cement the attachment bond between mother and child but also lay the foundation for the concordance of multiple brains, thereby creating a neural template for the consolidation of individuals into social groups.

Our findings contribute to the literature on cross-brain synchrony by showing a widespread cross-brain concordance to non-verbal stimuli that present no overt narrative. Most previous studies focused on the unfolding of stories with a clear narrative (Hasson et al., 2008; Wilson et al., 2008; Nguyen et al., 2019; Redcay and Moraczewski, 2020) and only a few presented stimuli with no verbal narrative, such as dance performance, music, or action observation (Herbec et al., 2015; Kostorz et al., 2020; Sachs et al., 2020). Stimuli of the mother-infant bond are fundamental and are immediately understood across cultures with no need for words, and the widespread neural concordance found here suggests that human cross-brain similarity does not rely solely on higher-order mechanisms of shared mentalization, language, and cognition. Our study demonstrates that in the context of the mother-infant bond, there is no need for a complex and elaborated narrative to synchronize the perceivers’ brain responses across widely-distributed brain networks.

Our findings indicate that when exposed to *Social* mother-infant stimuli, the participants’ brains synchronized more intensely than in response to the *Alone* contexts. Regions displaying cross-brains concordance to both *Social* and *Alone* attachment stimuli spanned from the occipital cortex to the prefrontal cortex, through temporal, limbic, and paralimbic regions. This is consistent with previous studies that measured cross-brain synchrony to naturalistic videos and showed high and stable ISC across extensive brain areas; from occipital regions including the visual cortex, fusiform gyrus, and the precuneus, to frontal regions including the OFC and temporal regions, and paralimbic regions, including the insula (Kauppi et al., 2010; Gao et al., 2020). Here, the broad-band inter-individual correlated response included regions of the caregiving network as well as other subcortical regions. This may be attributed to the universal nature of the maternal-infant bond and its evolutionary significance, which not only recruits substantial resources but also glues mothers into a uniform neural response. Our stimuli comprising daily ecological contexts of mothers and infants were highly relevant to the participants, who were at the stage of forming an exclusive and meaningful bond with their infant and allocating their physical, mental, and emotional energies to this survival-related task (Tronick, 1989; Feldman, 2020). Such bonding process involves neural reorganization of the PCN, which becomes highly sensitive to social cues (Kim, 2016). Our widespread findings are also consistent with research showing that ISC increases when humans share similar interests or psychological perspective (Lahnakoski et al., 2014; Hyon et al., 2020) or when they empathize with the stimuli (Borja Jimenez et al., 2020), indicating that similarity of mental focus, perspective-taking, and empathy are reflected in increased neural resemblance.

Both the top-down and bottom-up analyses illuminated the involvement of the PCN in the convergent cross-brain processing of the stimuli. These findings highlight the centrality of the PCN from a new angle; the network underpinning the formation of affiliative bonds that activates to attachment-related cues in parents (Abraham et al., 2014; Atzil et al., 2014), children (Pratt et al., 2018; Ulmer-Yaniv et al., 2022), romantic couples (Acevedo et al., 2012; Scheele et al., 2013), and close friends (Güroǧlu et al., 2008; Platek and Kemp, 2009; Parkinson et al., 2018) also functions to bind humans’ brains into a synchronized response and enhance neural resemblance among participants. The PCN includes conserved subcortical areas implicated in mammalian maternal care and these are connected via multiple ascending and descending projections to insular, temporal, and frontal regions (Rilling and Young, 2014; Feldman, 2015) and cohere to sustain human attachment, foster social motivation, and support higher-order sociocognitive processes, such as empathy and mentalization that enable the formation of human relationships. Our findings highlight the openness of this network to cross-brain processes in the presence of attachment-related cues.

Greater synchronization emerged in response to the *social* compared to the *alone* contexts in regions of the parenting network, such as the insula, PHG, ACC, STG, PFC, and striatum. Imaging studies that exposed parents to photos or videos of an unfamiliar infant report activations in the PFC, insula, ACC, PHG, and amygdala (Bartels and Zeki, 2004; Leibenluft et al., 2004; Nitschke et al., 2004; Strathearn et al., 2009; Landi et al., 2011). These regions were also selectively activated when parents observed videos of mother-infant social interactions and both activations and connectivity in these structures were linked with the degree of behavioral synchrony (Atzil et al., 2014; Abraham et al., 2017; Shimon-Raz et al., 2021; Ulmer-Yaniv et al., 2022). Of note, regions showing a main effect for the *Social* context also overlap with the areas reported in a meta-analytic map of the term “social” that tested response to social processes across 1302 studies (Fig. 3B, 3–4) (Yarkoni et al., 2011), indicating that these social areas not only activate to social stimuli but also show greater convergence when attachment-related social stimuli are presented. Our findings on the higher cross-brain similarity to the social contexts suggest that two-person representations of attachment generate lower variability of neural response across participants compared to single-person reminders. Such prototypic representation of the mother-infant attachment, an icon that stands at the heart of Western history as a symbol of utmost devotion, is internalized as the basic building-block of human sociality (Bowlby, 1969). Notably, auditory, visual, and somatosensory low-level regions also showed greater synchrony to the *Social* context. This may relate to the repetitive multi-modal nature of mother-infant interactions or from top-down circuitry based on the salience of these stimuli, but this hypothesis requires further research. The neural similarity across participants to mother-infant stimuli may also be relevant to the mechanism of “alomothering", the rearing of infants by adults other than the biological mother, a common practice across primate species and non-Western societies (Abraham & Feldman, 2018). The synchronized neural response to infant cues may charts the mechanism that enable multiple females in the clan to provide appropriate infant care, supporting that practice of allomothering that has served a key survival function across primate evolution.

The co-variability of ISC with shifts in behavioral synchrony is among the interesting and unique findings of our study. We found that fluctuations in the activity of the preregistered ACC tracked moment-by-moment variations in mother-infant synchrony, presenting the first evidence that links cross-brain synchrony with fluctuations in behavioral synchrony of the presented stimulus. The ACC is densely connected with sensory, limbic, and paralimbic regions (Etkin et al., 2006, 2011), underpins sensation, affective behavior, and decision-making (Peterson et al., 1999; Bush et al., 2000; Pavlvlović et al., 2009), and supports self-relational processes (Northoff et al., 2006; Ulmer-Yaniv et al., 2022). The ACC is also implicated in social observational learning in animals and humans (Burgos-Robles 2019), including observational fear learning in rodents (Allsop et al., 2018) and social decision-making in monkeys (Chang nature neuroscience 2018). Furthermore, the ACC has been shown to play a key role in brain-behavior linkage in the context of caregiving. For instance, the degree of mother-infant behavioral synchrony during free play has been associated with her ACC response to the presentation of synchronous versus asynchronous unfamiliar mother-infant interaction (Atzil et al., 2014). Similarly, mothers’ ACC response to her own infant video, compared to unfamiliar infant, correlated with her behavioral synchrony observed in the home environment (Abraham et al., 2014). Moreover, young adults viewing videos of their own interactions with their mother from infancy, childhood, and adulthood, compared to unfamiliar mother-child videos, not only showed greater ACC response to self-videos but also increased functional connectivity between ACC and insula in response to attachment reminders (Ulmer-Yaniv et al., 2022), highlighting the ACC-insula interface in the integration of interceptive and exteroceptive inputs in the context of attachment (Dosenbach et al., 2007; Medford and Critchley, 2010). Here we add to the ACC’s role in the consolidation of attachment representations also the integrative function of synchronizing activity across multiple brains to attachment cues in a manner that binds all brain to the presented stimuli.

While our first two pre-registered hypotheses were supported by the findings, the third hypothesis, which postulated greater cross-brain synchrony under OT, was not. OT has been repeatedly linked with parental caregiving behaviors (Olazábal and Young, 2006; Levine et al., 2007; Galbally et al., 2011) and OT administration modified activations in the PCN to attachment stimuli (Riem et al., 2011a; Owen et al., 2013; Wu et al., 2022). However, unlike levels of activation, no differences emerged in cross-brain correlations to attachment stimuli under OT versus PBO in both the social and alone contexts. While prior studies showed that OT modulated functional connectivity within- and between-networks, such as the DMN and salience network (Jiang et al., 2021; Zheng et al., 2021; Wu et al., 2022), cross-person correspondence to attachment cues has not been studied.

Possibly, attachment-related stimuli elicit such broad-band cross-brain resemblance in post-partum mothers that this may have created a ceiling effect. Overall, our results are consistent with the consensus in the field that OT effects are person-, time-, and context-sensitive (Bartz et al., 2010).

Limitations of the study mainly concern the specificity of our sample. As this is the first study of cross-brain synchrony to attachment stimuli, we chose to test mothers for whom such stimuli are the most relevant, but future studies are needed to assess cross-brain synchrony in fathers and non-parents to attachment cues. We believe that representations of the mother-infant bond would unify the neural response in all adult members of the species, but such hypothesis requires further research. Further research is also needed to assess whether psychopathologies involving social dysfunction, such as autism or depression, which have been associated with diminished cross-brain synchrony to a variety of stimuli (Salmi et al., 2013; Komulainen et al., 2021) also exhibit attenuated ISC to infant stimuli, and particularly relevant in this context are mothers suffering from postpartum depression. It would be of interest to see cross-brain synchrony to the father-infant bond in mothers, fathers, and non-parents and whether and how such synchrony differs from the current results of the mother-infant relationship. Finally, it is important to study neural response to negatively-valance attachment cues, such as infant cry or distress and to assess whether they trigger similar cross-individual neural resemblance as do positive moments of mother-infant interactions. Finally, as we preferred using real-life naturalistic stimuli, our stimuli are not fully controlled for low level properties, a condition that is generally an issue in ecological brain research.

To our knowledge, this is the first study to show neural synchronization across multiple brains in response to attachment stimuli. We showed, utilizing both theory-driven and data-driven analytic approaches, a widespread cross-person neural synchrony to attachment reminders, with greater synchrony in response to mother-infant social cues. Cross-person correlations in the ACC dynamically tracked moment-by-moment variations in mother-infant synchrony, suggesting that moments of coordinated behavior dynamically trigger uniformity across brains. Taken together, our findings highlight the primary attachment as a core survival-related phenomenon that recruits substantial neural resources and suggest that attachment bonds may provide a neural template for the consolidation of humans into social groups.

## Acknowledgments

The study was supported by The Simms/Mann Chair to Ruth Feldman and by the Bezos Family Foundation.

