## Supplementary for "Attachment Stimuli Trigger Widespread Synchrony across Multiple Brains"

**Extended data**

**Figure1-1**. 2*×*2 Bayesian repeated measures ANOVA results (*Infant* /*Mother alone* video *×* *PBO-OT*) in the PCN.

|  | **BF_10_** | **BF_incl_** |
| --- | --- | --- |
| *Mother alone/Infant alone* main effect | 0.24 | 0.16 |
| *PBO-OT* main effect | 0.22 | 0.18 |
| *Mother alone/Infant alone × PBO-OT* interaction | 0.05 | 0.07 |

2*×*2 Bayesian repeated measures ANOVA results (*Infant*/*Mother alone* video*×* *PBO-OT*) demonstrated evidence for the lack of difference between the 2 *Alone* context videos ISCs, between *PBO* and *OT* and the lack of interaction effect between them, through the PCN. PCN, Parental Caregiving Network.

**Figure1-2**. 2*×*2 Bayesian repeated measures ANOVA results (*Free-play*/*Breastfeeding* video *×* *PBO-OT*) in the PCN

|  | **BF_10_** | **BF_incl_** |
| --- | --- | --- |
| *Breastfeeding/Free-play main effect* | 0.34 | 0.79 |
| *PBO-OT* main effect | 0.24 | 0.73 |
| *Free-play*/*Breastfeeding × PBO-OT* interaction | 0.08 | 0.98 |

2*×*2 Bayesian repeated measures ANOVA results (*Free-play*/*Breastfeeding* video *×* *PBO-OT*) revealed the lack of difference between the 2 *Social* context videos ISCs between *PBO* and *OT* and the lack of interaction effect between them, through the Parental Caregiving Network (PCN).

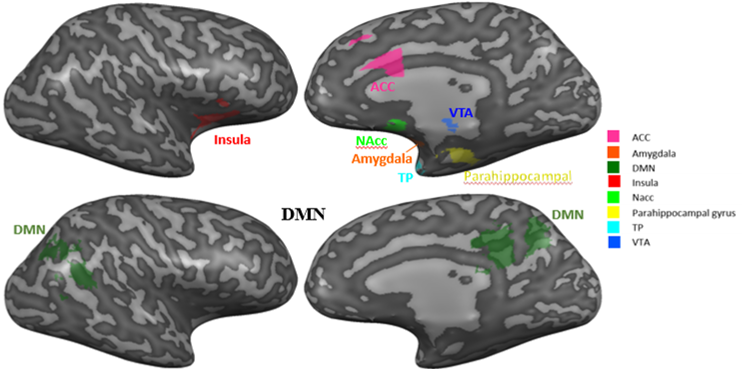

**Figure 1-3.** Eight preregistered ROIs including the ACC, amygdala, insula, NAcc, PHG, TP, VTA in the PCN (upper) and the DMN (footer). ACC, anterior cingulate cortex; NAcc, nucleus accumbens, PHG, parahippocampal gyrus; TP, temporal pole; VTA, ventral tegmental area, PCN, parental caregiving network; DMN, default mode network.

|  | **Infant** | | | **Mother** | | |
| --- | --- | --- | --- | --- | --- | --- |
| **behavioral synchrony level** | **Vocalization** | **Gaze** | **Affect** | **Vocalization** | **Gaze** | **Affect** |
| **1** | no vocalization | aversion | neutral | Motherese | to infant's face | neutral |
| **2** | no vocalization | to environment/object | neutral | Motherese | to infant's face | neutral |
| **3** | no vocalization | to mother | neutral | Motherese | to infant's face | neutral |
| **4** | no vocalization | to environment/object | positive (smile) | Motherese | to infant's body | neutral |
| **5** | no vocalization | to mother | neutral | Motherese | to infant's face | positive (smile) |
| **6** | no vocalization | to mother | positive (smile) | Motherese | to infant's body | neutral |
| **7** | no vocalization | to mother | positive (smile) | Motherese | to infant's face | neutral |
| **8** | cooing | to mother | positive (smile) | Motherese | to infant's face | neutral |
| **9** | no vocalization | to mother | positive (smile) | Motherese | to infant's body | positive (smile) |
| **10** | no vocalization | to environment/object | positive (smile) | Motherese | to infant's face | positive (smile) |
| **11** | no vocalization | to mother | positive (smile) | Motherese | to infant's face | positive (smile) |
| **12** | cooing | to environment/object | positive (smile) | Motherese | to infant's body | positive (smile) |
| **13** | cooing | to mother | positive (smile) | Motherese | to infant's body | positive (smile) |
| **14** | cooing | to mother | positive (smile) | Motherese | to infant's face | positive (smile) |
| **15** | laugh | to mother | positive (smile) | Motherese | to infant's face | positive (smile) |

**Figure 1-4**. Behavioral coding of mother-infant synchronization during free play interaction

Mothe and infant behavior during the free-play interaction was coded over 3 non-verbal categories: gaze, affect, and vocalization. The synchronization level indicates the degree of compatibility between the mother and the baby in three categories, with a higher score indicating a higher level of behavioral synchronization.

**Figure 2-2.** ISC threshold values for theory driven analysis (ROIs)

|  |  | Threshold | TP | ACC* | Insula* | PHG* | Amygdala | NAcc* | VTA | DMN* |
| --- | --- | --- | --- | --- | --- | --- | --- | --- | --- | --- |
| *PBO*  ISC (*p*) | Alone | 0.12 | 0.0187  (0.311) | 0.0556  (0.053) | 0.0607  (0.027) | 0.0182  (0.235) | 0.0870  (0.002) | 0.1688  (0.001) | -0.0145  (0.659) | 0.1494  (0.001) |
|  | Social | 0.13 | 0.0730  (0.022) | 0.1079  (0.002) | 0.1308  (0.001) | 0.1394  (0.001) | 0.0343  (0.128) | 0.1340  (0.001) | -0.0179  (0.675) | 0.1453  (0.001) |
| *OT*  ISC (*p*) | Alone | 0.12 | -0.0251  (0.795) | 0.1031  (0.004) | 0.0726  (0.008) | 0.0228  (0.267) | 0.0075  (0.716) | 0.1237  (0.001) | 0.0693  (0.005) | 0.0726  (0.001) |
|  | Social | 0.11 | 0.0860  (0.007) | 0.1656  (0.001) | 0.1665  (0.001) | 0.1074  (0.001) | 0.0296  (0.102) | 0.1606  (0.001) | 0.0481  (0.965) | 0.1665  (0.001) |

ISC, *p* and threshold values of 8 predefined ROIs for the *Alone* and *Social* conditions under PBO and OT. *, ROIs had significant above threshold ISC in at least one condition. All results are FDR corrected.

**Figure 2-3.** ISC threshold values for data driven analysis (parcels)

| Condition | | ISC |
| --- | --- | --- |
| PBO | Alone | 0.1379 |
|  | Social | 0.1426 |
| OT | Alone | 0.1368 |
|  | Social | 0.1413 |

| Parcel number | Anatomical area | ISC | |
| --- | --- | --- | --- |
| *Exclusive to Social context* | |  | |
| PBO |  |  | |
| 8 | Prefrontal ventrolateral lateral parietal | 0.15 | |
| 16 | Insula | 0.17 | |
| 33 | Motor cortex | 0.16 | |
| 96 | Hippocampal gyrus medial temporal | 0.15 | |
| 100 | Cerebellum | 0.15 | |
| 123 | Striatum ventral striatum | 0.16 | |
| 125 | Striatum ventral striatum | 0.17 | |
| 147 | Dorsolateral PFC | 0.17 | |
| 150 | ACC | 0.15 | |
| 162 | Motor cortex | 0.18 | |
| 165 | Premotor cortex | 0.16 | |
| 188 | Temporal pole | 0.19 | |
| 225 | Precuneus | 0.16 | |
| OT |  |  | |
| 4 | Orbitofrontal cortex | 0.182 | |
| 14 | Dorsolateral PFC | 0.147 | |
| 19 | Frontoparietal | 0.199 | |
| 30 | Prefrontal parietal | 0.152 | |
| 32 | Premotor cortex | 0.193 | |
| 39 | Somatosensory cortex | 0.182 | |
| 89 | Midline | 0.185 | |
| 91 | precuneus | 0.199 | |
| 137 | Orbitofrontal cortex | 0.166 | |
| 139 | Orbitofrontal cortex | 0.149 | |
| 142 | Prefrontal cortex | 0.147 | |
| 144 | Prefrontal cortex | 0.169 | |
| 166 | Premotor | 0.221 | |
| 169 | Putamen, striatum | 0.165 | |
| 229 | Hippocampus, hippocampal cortex |  | |
| 230 | Parahippocampal cortex | | 0.155 |
| 240 | Cerebellum | | 0.203 |
| 246 | Cerebellum | | 0.173 |
| 259 | Striatum accumbens | | 0.146 |

**Figure 2-1.** 118 parcels involved in stimulus processing ISCs

whole brain

| Parcel number | Anatomical area | ISC |
| --- | --- | --- |
| *Exclusive to Social context* | |  |
| PBO and OT |  |  |
| 27 PBO | Premotor | 0.168 |
| 27 OT | Premotor | 0.15 |
| 37 PBO | Insula | 0.154 |
| 37 OT | Insula | 0.192 |
| 53 PBO | Temporal pole, superior temporal | 0.198 |
| 53 OT | Temporal pole, superior temporal | 0.175 |
| 143 PBO | Prefrontal cortex | 0.184 |
| 143 OT | Prefrontal cortex | 0.152 |
| 155 PBO | Anterior insula | 0.143 |
| 155 OT | Anterior insula | 0.149 |
| 170 PBO | Insula | 0.162 |
| 170 OT | Insula | 0.185 |
| 173 PBO | Posterior insula | 0.170 |
| 173 OT | Posterior insula | 0.146 |
| 184 PBO | Parietal ipl | 0.169 |
| 184 OT | Parietal ipl | 0.164 |
| 192 PBO | Superior temporal | 0.250 |
| 192 OT | Superior temporal | 0.194 |
| 217 PBO | Paralimbic | 0.206 |
| 217 OT | Paralimbic | 0.163 |
| *Exclusive to Alone context* | |  |
| OT |  |  |
| 243 | Cerebellum | 0.153 |
| *Involved in Alone and Social contexts* | |  |
| PBO |  |  |
| 242 alone | Cerebellum | 0.161 |
| 242 social | Cerebellum | 0.152 |
| OT |  |  |
| 44 alone | Superior parietal | 0.160 |
| 44 social | Superior parietal | 0.171 |
| 86 alone | Posterior cingulate precuneus posterior | 0.160 |
| 86 social | Posterior cingulate precuneus posterior | 0.197 |
| 177 alone | Intraparietal sulcus | 0.142 |
| 177 social | Intraparietal sulcus | 0.199 |
| 178 alone | Parietal | 0.163 |
| 178 social | Parietal | 0.165 |
| 183 alone | Temporoparietal junction | 0.138 |
| 183 social | Temporoparietal junction | 0.145 |
| 208 alone | Occipital visual | 0.190 |
| 208 social | Occipital visual | 0.172 |
| 224 alone | Parietal junction, posterior cingulate | 0.164 |
| 224 social | Parietal junction, posterior cingulate | 0.159 |
| 226 alone | Sensorimotor | 0.152 |

| Parcel number | Anatomical area | | ISC |
| --- | --- | --- | --- |
| *Involved in Alone and Social contexts* | | |  |
| OT | |  |  |
| 226 social | | Sensorimotor | 0.196 |
| PBO&OT | |  |  |
| 22 PBO Alone | | Dorsolateral PFC | 0.217 |
| 22 PBO social | | Dorsolateral PFC | 0.175 |
| 22 OT alone | | Dorsolateral PFC | 0.174 |
| 22 OT social | | Dorsolateral PFC | 0.236 |
| 26 PBO alone | | Motor cortex sensorimotor | 0.156 |
| 26 PBO social | | Motor cortex sensorimotor | 0.155 |
| 26 OT alone | | Motor cortex sensorimotor | 0.150 |
| 31 PBO alone | | Temporal parietal | 0.158 |
| 31 PBO social | | Temporal parietal | 0.146 |
| 31 OT alone | | Temporal parietal | 0.162 |
| 31 OT social | | Temporal parietal | 0.195 |
| 38 PBO alone | | Premotor | 0.247 |
| 38 PBO social | | Premotor | 0.241 |
| 38 OT alone | | Premotor | 0.231 |
| 38 OT social | | Premotor | 0.257 |
| 41 PBO alone | | Premotor | 0.147 |
| 41 PBO social | | Premotor | 0.198 |
| 41 OT alone | | Premotor | 0.155 |
| 41 OT social | | Premotor | 0.249 |
| 42 PBO alone | | Precuneus | 0.158 |
| 42 PBO social | | Precuneus | 0.145 |
| 42 OT alone | | Precuneus | 0.187 |
| 42 OT social | | Precuneus | 0.185 |
| 43 PBO alone | | Parietal cortex | 0.181 |
| 43 PBO social | | Parietal cortex | 0.172 |
| 43 OT social | | Parietal cortex | 0.231 |
| 45 PBO alone | | Somatosensory | 0.143 |
| 45 PBO social | | Somatosensory | 0.168 |
| 45 OT alone | | Somatosensory | 0.186 |
| 45 OT social | | Somatosensory | 0.235 |
| 46 PBO alone | | Somatosensory | 0.301 |
| 46 PBO social | | Somatosensory | 0.344 |
| 46 OT alone | | Somatosensory | 0.287 |
| 46 OT social | | Somatosensory | 0.341 |
| 47 PBO alone | | Inferior parietal | 0.138 |
| 47 PBO social | | Inferior parietal | 0.151 |
| 47 OT social | | Inferior parietal | 0.150 |
| 48 PBO alone | | Default mode** | 0.197 |
| 48 PBO social | | Default mode** | 0.166 |
| 48 OT alone | | Default mode** | 0.157 |
| 48 OT social | | Default mode** | 0.161 |

| Parcel number | Anatomical area | ISC |
| --- | --- | --- |
| *Involved in Alone and Social contexts* | |  |
| PBO&OT |  |  |
| 49 PBO alone | Medial temporal | 0.252 |
| 49 PBO social | Medial temporal | 0.245 |
| 49 OT alone | Medial temporal | 0.284 |
| 49 OT social | Medial temporal | 0.328 |
| 50 PBO alone | Temporal sulcus | 0.302 |
| 50 PBO social | Temporal sulcus | 0.268 |
| 50 OT alone | Temporal sulcus | 0.279 |
| 50 OT social | Temporal sulcus | 0.279 |
| 54 PBO alone | Temporal sulcus superior temporal | 0.211 |
| 54 PBO social | Temporal sulcus superior temporal | 0.211 |
| 54 OT alone | Temporal sulcus superior temporal | 0.312 |
| 54 OT social | Temporal sulcus superior temporal | 0.196 |
| 61 PBO alone | primary auditory | 0.242 |
| 61 PBO social | primary auditory | 0.285 |
| 61 OT alone | primary auditory | 0.224 |
| 61 OT social | primary auditory | 0.256 |
| 62 PBO alone | posterior insula | 0.179 |
| 62 PBO social | posterior insula | 0.280 |
| 62 OT alone | posterior insula | 0.218 |
| 62 OT social | posterior insula | 0.270 |
| 63 PBO alone | Superior temporal | 0.241 |
| 63 PBO social | Superior temporal | 0.340 |
| 63 OT alone | Superior temporal | 0.225 |
| 63 OT social | Superior temporal | 0.227 |
| 64 PBO alone | Lateral temporal | 0.209 |
| 64 PBO social | Lateral temporal | 0.214 |
| 64 OT alone | Lateral temporal | 0.158 |
| 64 OT social | Lateral temporal | 0.176 |
| 65 PBO alone | Temporal sulcus | 0.352 |
| 65 PBO social | Temporal sulcus | 0.297 |
| 65 OT alone | Temporal sulcus | 0.294 |
| 65 OT social | Temporal sulcus | 0.283 |
| 66 PBO alone | Fusiform | 0.247 |
| 66 PBO social | Fusiform | 0.240 |
| 66 OT alone | Fusiform | 0.203 |
| 66 OT social | Fusiform | 0.245 |
| 67 PBO alone | Fusiform | 0.254 |
| 67 PBO social | Fusiform | 0.213 |
| 67 OT alone | Fusiform | 0.236 |
| 67 OT social | Fusiform | 0.251 |
| 68 PBO alone | Parahippocampal | 0.223 |
| 68 PBO social | Parahippocampal | 0.238 |
| 68 OT alone | Parahippocampal | 0.276 |
| 68 OT social | Parahippocampal | 0.282 |
| 69 PBO alone | Visual cortex | 0.224 |
| 69 PBO social | Visual cortex | 0.209 |
| 69 OT alone | Visual cortex | 0.197 |
| 69 OT social | Visual cortex | 0.270 |

| Parcel number | Anatomical area | ISC |
| --- | --- | --- |
| *Involved in Alone and Social contexts* | |  |
| PBO&OT |  |  |
| 71 PBO alone | Fusiform | 0.189 |
| 71 PBO social | Fusiform | 0.191 |
| 71 OT alone | Fusiform | 0.179 |
| 71 OT social | Fusiform | 0.216 |
| 72 PBO alone | Visual fusiform | 0.194 |
| 72 PBO social | Visual fusiform | 0.187 |
| 72 OT alone | Visual fusiform | 0.230 |
| 72 OT social | Visual fusiform | 0.240 |
| 73 PBO alone | Visual cortex occipital | 0.256 |
| 73 PBO social | Visual cortex occipital | 0.341 |
| 73 OT alone | Visual cortex occipital | 0.274 |
| 73 OT social | Visual cortex occipital | 0.403 |
| 74 PBO alone | Visual cortex occipital | 0.378 |
| 74 PBO social | Visual cortex occipital | 0.368 |
| 74 OT alone | Visual cortex occipital | 0.271 |
| 74 OT social | Visual cortex occipital | 0.432 |
| 75 PBO social | Occipital parietal | 0.220 |
| 75 OT alone | Occipital parietal | 0.161 |
| 75 OT social | Occipital parietal | 0.264 |
| 76 PBO alone | Visual occipital | 0.220 |
| 76 PBO social | Visual occipital | 0.246 |
| 76 OT alone | Visual occipital | 0.267 |
| 76 OT social | Visual occipital | 0.244 |
| 77 PBO alone | Cuneus | 0.222 |
| 77 PBO social | Cuneus | 0.181 |
| 77 OT alone | Cuneus | 0.220 |
| 77 OT social | Cuneus | 0.230 |
| 78 PBO alone | Occipital gyrus | 0.340 |
| 78 PBO social | Occipital gyrus | 0.362 |
| 78 OT alone | Occipital gyrus | 0.307 |
| 78 OT social | Occipital gyrus | 0.414 |
| 79 PBO alone | Visual cortex | 0.230 |
| 79 PBO social | Visual cortex | 0.179 |
| 79 OT alone | Visual cortex | 0.282 |
| 79 OT social | Visual cortex | 0.215 |
| 80 PBO alone | Occipital cuneus | 0.221 |
| 80 PBO social | Occipital cuneus | 0.198 |
| 80 OT alone | Occipital cuneus | 0.277 |
| 80 OT social | Occipital cuneus | 0.214 |
| 81 PBO alone | Inferior occipital | 0.306 |
| 81 PBO social | Inferior occipital | 0.245 |
| 81 OT alone | Inferior occipital | 0.278 |
| 81 OT social | Inferior occipital | 0.302 |
| 82 PBO alone | Parietal network visual | 0.216 |
| 82 PBO social | Parietal network visual | 0.152 |
| 82 OT alone | Parietal network visual | 0.221 |
| 82 OT social | Parietal network visual | 0.164 |

| Parcel number | Anatomical area | | ISC |
| --- | --- | --- | --- |
| *Involved in Alone and Social contexts* | | |  |
| PBO&OT | |  |  |
| 87 PBO alone | | Accumbens | 0.148 |
| 87 OT social | | Accumbens | 0.155 |
| 90 PBO alone | | Precuneus | 0.147 |
| 90 PBO social | | Precuneus | 0.153 |
| 90 OT alone | | Precuneus | 0.141 |
| 90 OT social | | Precuneus | 0.170 |
| 95 PBO alone | | Hippocampus | 0.141 |
| 95 OT social | | Hippocampus | 0.196 |
| 98 PBO social | | Hippocampus | 0.148 |
| 98 OT alone | | Hippocampus | 0.155 |
| 163 PBO social | | Somatosensory cortices | 0.266 |
| 163 OT alone | | Somatosensory cortices | 0.175 |
| 163 OT social | | Somatosensory cortices | 0.227 |
| 166 PBO alone | | Premotor | 0.211 |
| 166 PBO social | | Premotor | 0.208 |
| 166 OT alone | | Premotor | 0.190 |
| 166 OT social | | Premotor | 0.221 |
| 171 PBO alone | | Motor cortex | 0.147 |
| 171 PBO social | | Motor cortex | 0.237 |
| 171 OT alone | | Motor cortex | 0.141 |
| 171 OT social | | Motor cortex | 0.232 |
| 175 PBO alone | | Premotor | 0.203 |
| 175 PBO social | | Premotor | 0.195 |
| 175 OT alone | | Premotor | 0.182 |
| 175 OT social | | Premotor | 0.253 |
| 176 PBO alone | | Precuneus | 0.158 |
| 176 PBO social | | Precuneus | 0.162 |
| 176 OT alone | | Precuneus | 0.155 |
| 176 OT social | | Precuneus | 0.183 |
| 179 PBO alone | | Intraparietal premotor | 0.217 |
| 179 PBO social | | Intraparietal premotor | 0.253 |
| 179 OT alone | | Intraparietal premotor | 0.204 |
| 179 OT social | | Intraparietal premotor | 0.273 |
| 180 PBO alone | | Auditory cortex | 0.279 |
| 180 PBO social | | Auditory cortex | 0.368 |
| 180 OT alone | | Auditory cortex | 0.295 |
| 180 OT social | | Auditory cortex | 0.354 |
| 181 PBO alone | | Somatosensory cortex | 0.155 |
| 181 PBO social | | Somatosensory cortex | 0.278 |
| 181 OT alone | | Somatosensory cortex | 0.179 |
| 181 OT social | | Somatosensory cortex | 0.293 |

| Parcel number | Anatomical area | | ISC |
| --- | --- | --- | --- |
| *Involved in Alone and Social conditions* | | |  |
| 182 PBO social | | Angular gyrus | 0.169 |
| 182 OT alone | | Angular gyrus | 0.139 |
| 191 PBO alone | | Auditory cortex | 0.206 |
| 191 PBO social | | Auditory cortex | 0.352 |
| 191 OT alone | | Auditory cortex | 0.242 |
| 191 OT social | | Auditory cortex | 0.297 |
| 197 PBO alone | | Auditory cortex superior temporal | 0.158 |
| 197 PBO social | | Auditory cortex superior temporal | 0.273 |
| 197 OT social | | Auditory cortex superior temporal | 0.161 |
| 198 PBO alone | | Parahippocampal | 0.153 |
| 198 OT alone | | Parahippocampal | 0.220 |
| 198 OT social | | Parahippocampal | 0.202 |
| 200 PBO alone | | Fusiform | 0.184 |
| 200 PBO social | | Fusiform | 0.188 |
| 200 OT alone | | Fusiform | 0.227 |
| 200 OT social | | Fusiform | 0.204 |
| 203 PBO alone | | Visual motion default mode | 0.230 |
| 203 PBO social | | Visual motion default mode | 0.185 |
| 203 OT alone | | Visual motion default mode | 0.284 |
| 203 OT social | | Visual motion default mode | 0.225 |
| 204 PBO alone | | Occipital cortex | 0.216 |
| 204 PBO social | | Occipital cortex | 0.277 |
| 204 OT alone | | Occipital cortex | 0.270 |
| 204 OT social | | Occipital cortex | 0.299 |
| 205 PBO alone | | Parahippocampal cortex | 0.163 |
| 205 PBO social | | Parahippocampal cortex | 0.149 |
| 205 OT alone | | Parahippocampal cortex | 0.139 |
| 205 OT social | | Parahippocampal cortex | 0.194 |
| 206 PBO alone | | Visual occipital fusiform | 0.254 |
| 206 PBO social | | Visual occipital fusiform | 0.199 |
| 206 OT alone | | Visual occipital fusiform | 0.249 |
| 206 OT social | | Visual occipital fusiform | 0.238 |
| 207 PBO alone | | Fusiform | 0.159 |
| 207 PBO social | | Fusiform | 0.145 |
| 207 OT alone | | Fusiform | 0.219 |
| 207 OT social | | Fusiform | 0.150 |
| 209 PBO alone | | Visual | 0.294 |
| 209 PBO social | | Visual | 0.350 |
| 209 OT alone | | Visual | 0.279 |
| 209 OT social | | Visual | 0.364 |
| 210 PBO alone | | Visual occipital fusiform | 0.289 |
| 210 PBO social | | Visual occipital fusiform | 0.273 |
| 210 OT alone | | Visual occipital fusiform | 0.309 |
| 210 OT social | | Visual occipital fusiform | 0.286 |

| Parcel number | Anatomical area | | ISC |
| --- | --- | --- | --- |
| *Involved in Alone and Social conditions* | | |  |
| 211 PBO alone | | Early visual | 0.187 |
| 211 PBO social | | Early visual | 0.180 |
| 211 OT alone | | Early visual | 0.231 |
| 211 OT social | | Early visual | 0.219 |
| 212 PBO alone | | Visual occipital | 0.227 |
| 212 PBO social | | Visual occipital | 0.207 |
| 212 OT alone | | Visual occipital | 0.253 |
| 212 OT social | | Visual occipital | 0.272 |
| 213 PBO alone | | Visual | 0.159 |
| 213 PBO social | | Visual | 0.162 |
| 213 OT alone | | Visual | 0.208 |
| 213 OT social | | Visual | 0.180 |
| 214 PBO alone | | Inferior occipital | 0.264 |
| 214 PBO social | | Inferior occipital | 0.242 |
| 214 OT alone | | Inferior occipital | 0.252 |
| 214 OT social | | Inferior occipital | 0.298 |
| 215 PBO alone | | Cuneus | 0.218 |
| 215 PBO social | | Cuneus | 0.175 |
| 215 OT alone | | Cuneus | 0.236 |
| 215 OT social | | Cuneus | 0.185 |
| 216 PBO alone | | Early visual | 0.174 |
| 216 PBO social | | Early visual | 0.192 |
| 216 OT alone | | Early visual | 0.158 |
| 216 OT social | | Early visual | 0.197 |
| 247 PBO alone | | Cerebellum | 0.167 |
| 247 OT social | | Cerebellum | 0.144 |

**Figure 3-1.** 4×2×2 repeated measures ANOVA (*ROI* × *Context* × *PBO-OT)* effects on ISC in PCN ROIs.

|  | df | F score | P | Eta^2^ |
| --- | --- | --- | --- | --- |
| ROI main effect | 2.53,58.40 | 5.21 | 0.003 | 0.185 |
| *Context* main effect | 1,23 | 10.96 | 0.003 | 0.323 |
| *PBO-OT* main effect | 1,23 | 0.28 | 0.604 | 0.012 |
| *ROI × Context* interaction | 2.19, 50.52 | 2.70 | 0.072 | 0.105 |
| *ROI × PBO-OT* interaction | 2.45, 56.24 | 1.08 | 0.358 | 0.045 |
| *Context × PBO-OT* interaction | 1,23 | 0.13 | 0.724 | 0.006 |
| *ROI × Context × PBO-OT* interaction | 2.26, 51.93 | 0.57 | 0.589 | 0.024 |

In the table results of 3 factors repeated measures ANOVA (*ROI* × *Context* × *PBO-OT*) conducted in the Insula, ACC, NAcc, and the PHG. All results are Greenhouse-Geisser corrected. OT, oxytocin; PBO, placebo.

**Figure 3-2.** 2×2 repeated measures ANOVA (*Context* × *PBO- OT*) for each of the preregistered ROIs above ISC threshold.

|  | Insula | ACC | PHG | NAcc | DMN |
| --- | --- | --- | --- | --- | --- |
| *Context main effect, df (1,23)* | | | | | |
| F score | 14.55 | 3.70 | 11.064 | 0.00 | 0.07 |
| P | 0.00** | 0.07 | 0.00** | 0.97 | 0.80 |
| Eta^2^ | 0.39 | 014 | 0.33 | 0.00 | 0.00 |
| *PBO-OT main effect, df (1,23)* | | | | | |
| F score | 0.56 | 2.64 | 0.13 | 0.05 | 0.30 |
| P | 0.46 | 0.12 | 0.72 | 0.83 | 0.60 |
| Eta^2^ | 0.02 | 0.10 | 0.01 | 0.00 | 0.01 |
| *Context × PBO-OT interaction, df (1,23)* | | | | | |
| F score | 0.10 | 0.02 | 0.46 | 0.78 | 0.03 |
| P | 0.75 | 0.88 | 0.50 | 0.39 | 0.87 |
| Eta^2^ | 0.00 | 0.00 | 0.00 | 0.03 | 0.00 |

In the table are results of *Context* and *PBO-OT* main effects and *Context* × *PBO-OT* interaction in 5 ROIs involved in stimulus processing. Only the insula and the PHG showed significant main effect for *Context*. All results are Greenhouse-Geisser corrected. OT, oxytocin; PBO, placebo; ACC, anterior cingulate cortex; NAcc, nucleus accumbens; PHG, parahippocampal gyrus; DMN, Default mode network. **, p<.001.

| Parcel number | Anatomical area | F (1,23) | | p | | Cluster size | Cluster peak voxel | | |
| --- | --- | --- | --- | --- | --- | --- | --- | --- | --- |
|  |  |  |  |  |  |  | X | Y | Z |
| *Context* main effect | | |  | |  |  |  |  |  |
| 4 | R orbitofrontal cortex | 7.75 | | 0.0065 | | 2408 | 16 | 35 | -22 |
| 27 | R premotor cortex | 38.9 | | <0.0001 | | 4824 | 50 | -4 | 48 |
| 33 | R Motor cortex | 13.99 | | 0.00031 | | 6952 | 42 | -23 | 54 |
| 39 | Somatosensory | 11.90 | | 0.00084 | | 6704 | 21 | -33 | 70 |
| 53 | R Temporal pole | 20.40 | | <0.0001 | | 6856 | 53 | -11 | -21 |
| 54 | R Superior temporal | 8.53 | | 0.0043 | | 2712 | 51 | -33 | 0 |
| 73 | R Visual cortex | 8.89 | | 0.0036 | | 4992 | 31 | -83 | 21 |
| 75 | R Occipital parietal cortex | 8.17 | | 0.0052 | | 6992 | 19 | -81 | 42 |
| 89 | R Midline | 10.12 | | 0.0019 | | 3360 | 9 | -22 | 45 |
| 123 | R Striatum | 7.13 | | 0.0089 | | 3624 | 13 | 21 | 0 |
| 125 | R Ventral striatum | 7.72 | | 0.0066 | | 4992 | 15 | 9 | -9 |
| 142 | L Prefrontal cortex | 7.81 | | 0.0063 | | 4088 | -29 | 55 | 3 |
| 143 | L Prefrontal cortex | 15.43 | | 0.00016 | | 4592 | -42 | 48 | -6 |
| 147 | L Dorsolateral prefrontal cortex | 15.04 | | 0.00019 | | 7040 | -46 | 29 | 27 |
| 150 | L Anterior cingulate cortex | 7.68 | | 0.0067 | | 4576 | -4 | 18 | 47 |
| 155 | L Insula | 17.17 | | <0.0001 | | 4496 | -32 | 23 | 6 |
| 163 | L Primary somatosensory cortex | 11.58 | | 0.0009 | | 7056 | -56 | -3 | 7 |
| 165 | L Premotor cortex | 6.91 | | 0.010 | | 5448 | -45 | 0 | 50 |
| 171 | L Motor cortex | 8.24 | | 0.0050 | | 6728 | -50 | -23 | 42 |
| 181 | L Somatosensory cortex | 13.19 | | 0.00046 | | 6840 | -59 | -25 | 22 |
| 184 | L Inferior [parietal](https://neurosynth.org/analyses/terms/parietal) cortex | 28.20 | | <0.0001 | | 10072 | -53 | -43 | 39 |
| 188 | L Temporal pole | 9.92 | | 0.0022 | | 5992 | -49 | 7 | -15 |
| 191 | L Auditory cortex | 7.60 | | 0.007 | | 6872 | -58 | -29 | 4 |
| 192 | L Superior temporal | 12.93 | | 0.00052 | | 7544 | -57 | -47 | 6 |
| 217 | L [Paralimbic](https://neurosynth.org/analyses/terms/paralimbic) | 17.48 | | <0.0001 | | 2368 | -23 | -41 | 20 |
| 230 | L Parahippocampal gyrus | 13.98 | | 0.00031 | | 2512 | -32 | -40 | -3 |
| 246 | L Cerebellum | 9.75 | | 0.0023 | | 6752 | -42 | -63 | -46 |

**Figure 3-3.** Parcels with significant *context* main effects results and their coordinates of activation peaks.

2×2 repeated measures ANOVA (*Context* × *PBO- OT*) was performed to each of the 118 parcels that were involved in stimulus processing. The table presents 27 parcels that had significant Context main effect, together with their anatomical location (defined by Neurosynth), F scores, p values, cluster sizes and Cluster peak voxel**.** Abbreviation: L, left; R, right.

**Figure 3-4.** Statistical meta-analytic brain activation map of the term “social” across 1302 studies. The map was obtained through Neurosynth. RH, Right hemisphere; LH, left hemisphere.

RH	LH

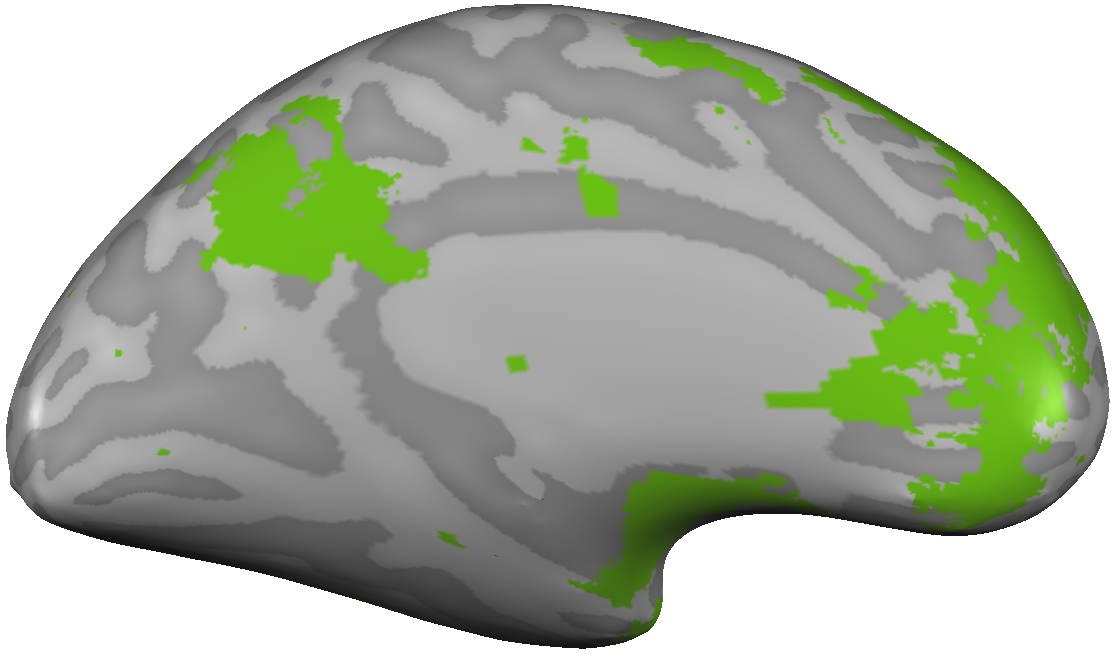

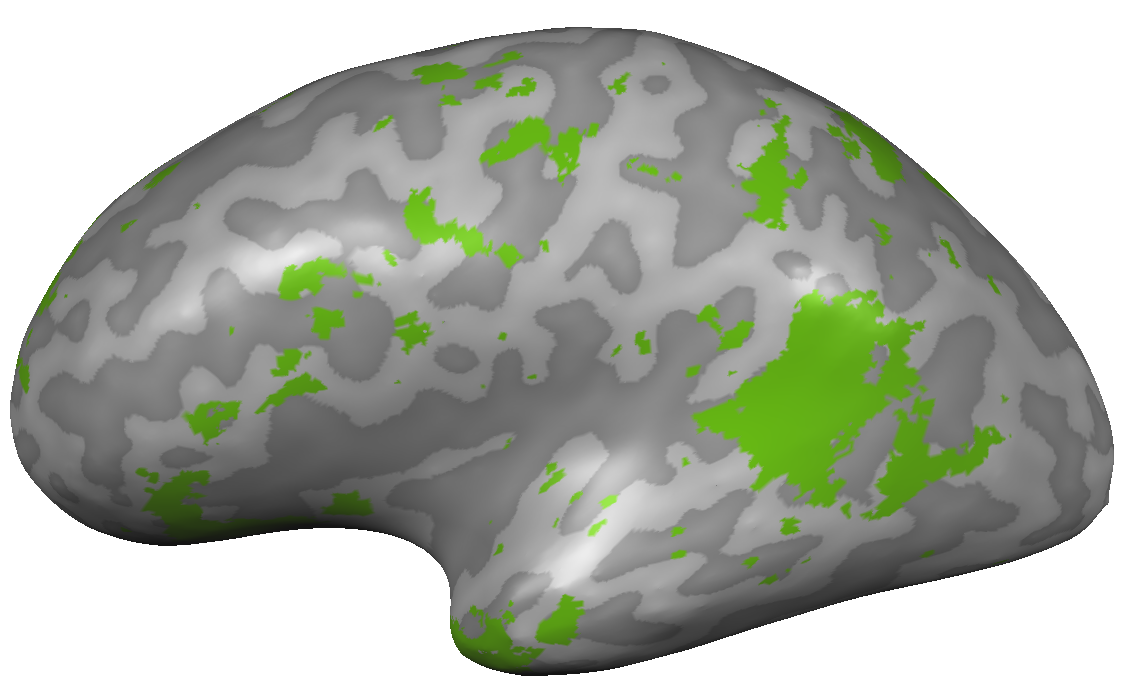

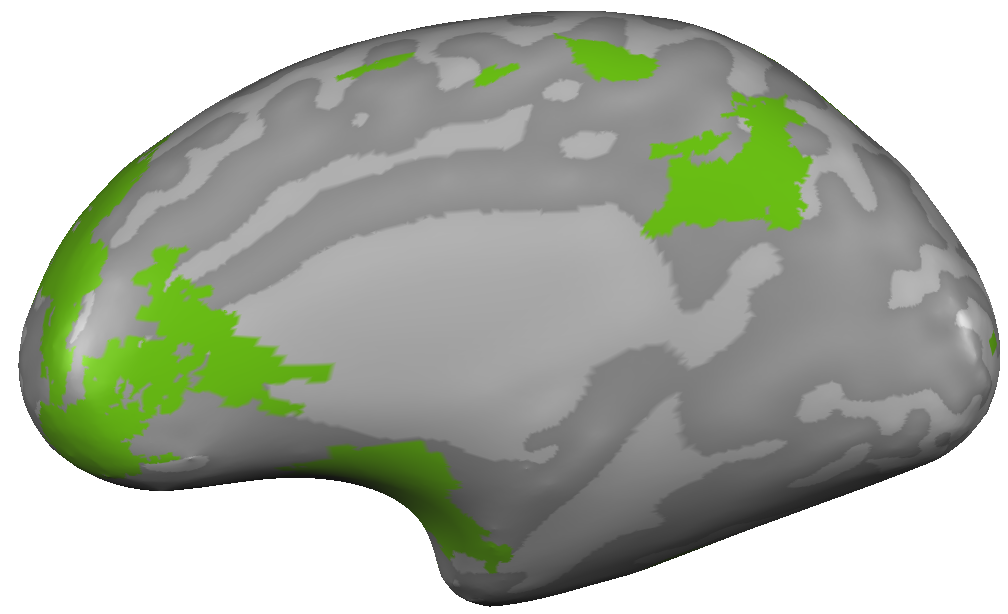

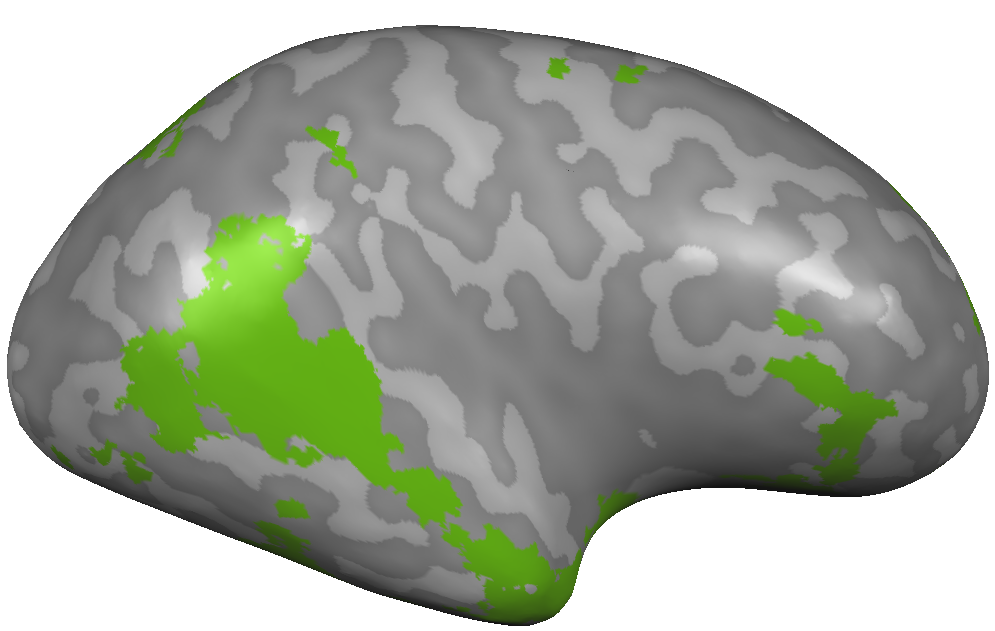

**Figure 3-5.** 2×2 repeated measures ANOVA (*Context* × *PBO- OT*) yielded a significant *Context* effect in 15 parcels in addition to PCN regions. These included the motor, somatosensory, visual, and orbitofrontal cortices. Their ISC and SE under the Alone (in blue) and the Social (in yellow) contexts are presented in the figure. Results are FDR corrected. OFC, Orbitofrontal cortex.

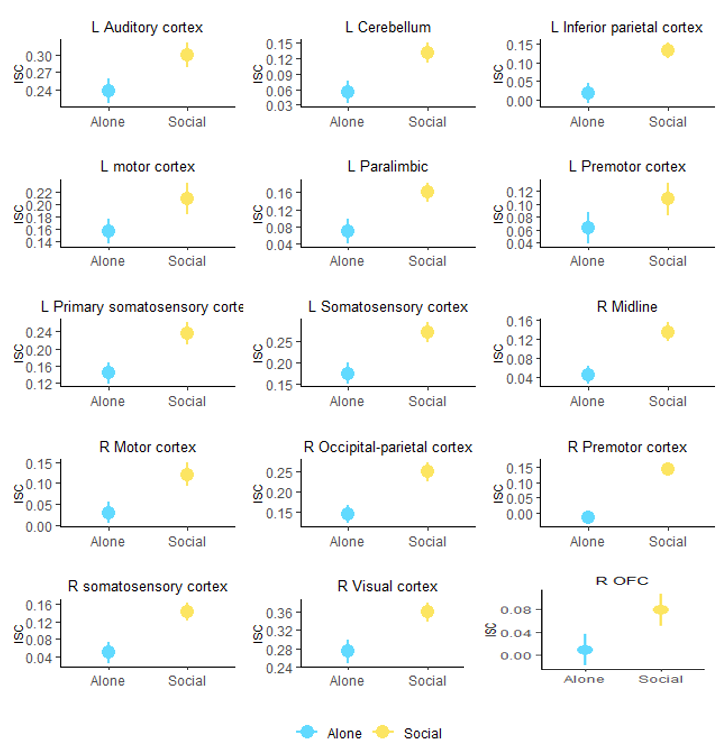

### **Figure 4-1.** Non-significant brain-behavior correlation results in ROIs.

|  | **Pearson's r** | **P** |
| --- | --- | --- |
| Insula | 0.32 | 0.054 |
| NAcc | 0.07 | 0.662 |
| PHG | 0.08 | 0.631 |
| DMN | 0.24 | 0.152 |

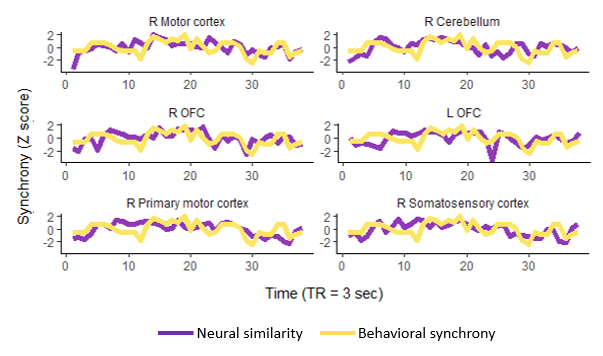

### **Figure 4-2**. Brain-behavior correlations results of data driven analyses. The figure depicts positive significant results of Pearson's correlations (not FDR corrected) between neural similarity (purple line) and mother-infant behavioral synchrony (yellow line) in the free-play video. Findings in the motor, primary motor, and somatosensory cortices, left and right OFC and in the cerebellum are presented. Abbreviations: OFC, orbitofrontal cortex; R, right; L, left.
